# A passive, camera-based head-tracking system for real-time, 3D estimate of head position and orientation in rodents

**DOI:** 10.1101/599365

**Authors:** Walter Vanzella, Natalia Grion, Daniele Bertolini, Andrea Perissinotto, Davide Zoccolan

## Abstract

Tracking head’s position and orientation of small mammals is crucial in many behavioral neurophysiology studies. Yet, full reconstruction of the head’s pose in 3D is a challenging problem that typically requires implanting custom headsets made of multiple LEDs or inertial units. These assemblies need to be powered in order to operate, thus preventing wireless experiments, and, while suitable to study navigation in large arenas, their application is unpractical in the narrow operant boxes employed in perceptual studies. Here we propose an alternative approach, based on passively imaging a 3D-printed structure, painted with a pattern of black dots over a white background. We show that this method is highly precise and accurate and we demonstrate that, given its minimal weight and encumbrance, it can be used to study how rodents sample sensory stimuli during a perceptual discrimination task and how hippocampal place cells represent head position over extremely small spatial scales.

## Introduction

Careful monitoring and quantification of motor behavior is essential to investigate a range of cognitive functions (such as motor control, active perception and spatial navigation) in a variety of different species. Examples include tracking eye movements in primate and non-primate species (Remmel, 1984; Stahl et al., 2000; Zoccolan et al., 2010; Kimmel et al., 2012; Wallace et al., 2013; Payne and Raymond, 2017), monitoring whisking activity in rodents (Knutsen et al., 2005; Perkon et al., 2011; Rigosa et al., 2017), and tracking the position of virtually any species displaying interesting navigation patterns – from bacteria (Berg and Brown, 1972) and invertebrate species (Mazzoni et al., 2005; Garcia-Perez et al., 2005; Mersch et al., 2013; Cavagna et al., 2017), to small terrestrial (Tort et al., 2006; Aragão et al., 2011) and aerial mammals (Tsoar et al., 2011; Yartsev and Ulanovsky, 2013) and birds (Attanasi et al., 2014). In particular, studies in laboratory animals aimed at measuring the neuronal correlates of a given behavior require tools that can accurately track it in time and space and record it along with the underlying neuronal signals.

A classical application of this approach is to track the position of a light-emitting diode (LED), mounted over the head of a rat or a mouse, while recording the activity of place cells in hippocampus (O’Keefe and Dostrovsky, 1971; O’Keefe and Nadel, 1978) or grid cells in entorhinal cortex (Fyhn et al., 2004; Hafting et al., 2005), with the goal of understanding how space is represented in these brain structures (Moser et al., 2008, 2015). It is also common, in studies of spatial representations, to track the *yaw* of the head (i.e., its orientation in the horizontal plane where the rodent navigates; see Figure 2C), to investigate the tuning of neurons in hippocampus, entorhinal cortex and other limbic structures for head direction, speed or angular velocity (Sargolini et al., 2006; Taube, 2007; Kropff et al., 2015; Acharya et al., 2016). In these applications, the yaw is tracked through an overhead video camera imaging two LEDs of different colors (e.g., red and green), mounted over the head stage of the neuronal recording system, and placed along the anteroposterior axis of the head. In some studies, this LED arrangement was also used to estimate the *pitch* of the head (i.e., its rotation about the interaural axis; see Figure 2C), by measuring the distance between the two LEDs in the image plane (Stackman and Taube, 1998; Bassett and Taube, 2001).

**Figure 1.**
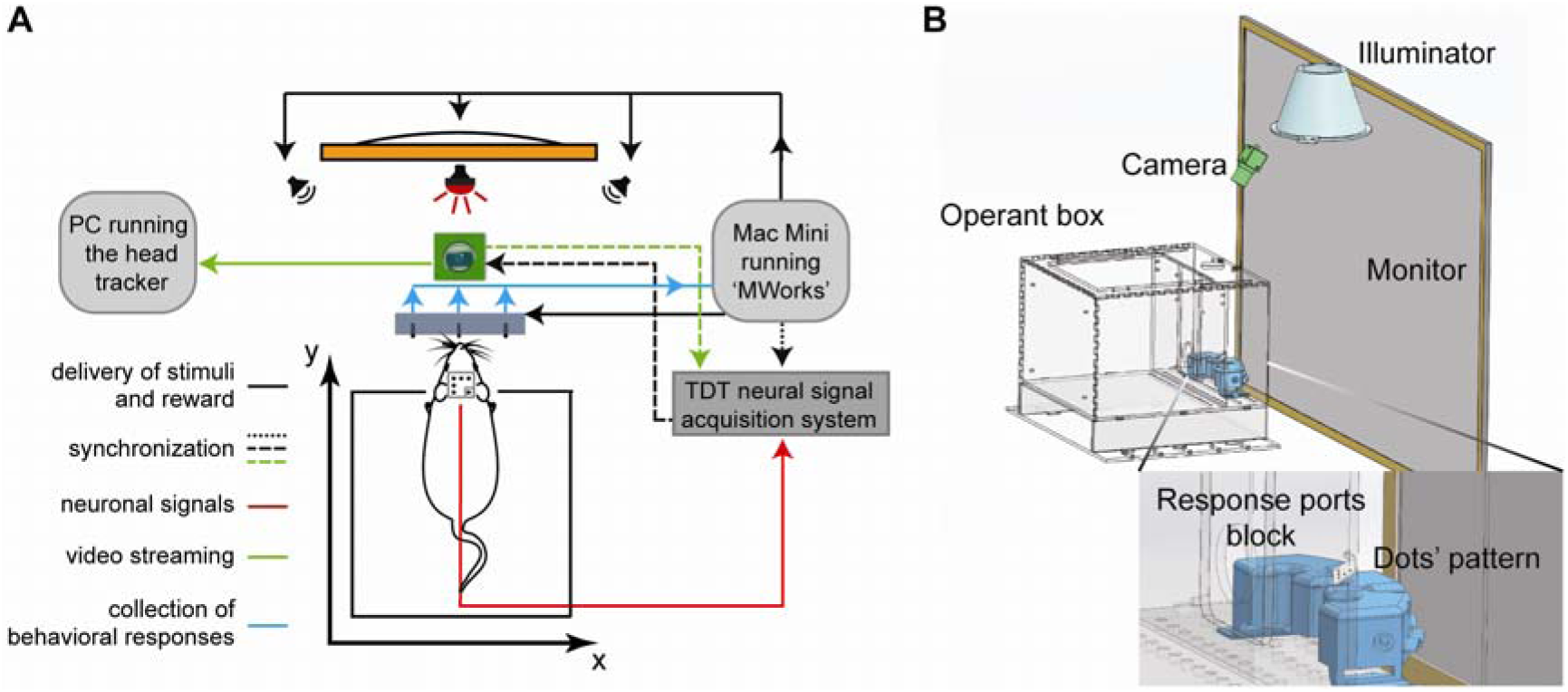
Illustration of the experimental rig where perceptual discrimination tests were combined with in-vivo neuronal recordings and real-time tracking of the head of a rat subject. (**A**) The animal was placed in an operant box and learned to protrude his head through an opening in one of the walls, so as to face the stimulus display (orange rectangle) and an array of response ports (gray rectangle), which also delivered liquid reward. Stimulus presentation and reward delivery (solid black arrows), as well as collection of the behavioral responses (solid cyan arrow) were controlled by the freeware application MWorks, running on a Mac mini (light-gray, rounded box on the right). MWorks also streamed unique codes (dotted black arrow) with the identity of the presented stimuli to the PC that controlled the recording of the neurophysiological signals from the rat brain (red arrow) through a Tucker-Davis Technologies (TDT) amplification/acquisition system (dark-gray box on the right). The head-tracking software run on a dedicated PC (light-gray, rounded box on the left), which received the video stream (solid green arrow) outputted by an industrial monochromatic CMOS camera (green box) placed above the operant box, along with a far-red light source (red bulb). The camera imaged a 3D pattern of dots mounted over the head of the animal (see Figure 2 for details). Image uptake was triggered by the TDT system (dashed black arrow), which, in turn, received from the camera a unique identification code for every acquired frame (dashed green arrow). The same code was also saved by the head-tracking software. The *x* and *y* axes (thick black arrows) indicate the Cartesian plane corresponding to the floor of the operant box. (**B**) A CAD rendering of some of the key elements of the experimental rig. The drawing allows appreciating the relative size and position of the operant box (light gray), stimulus display (dark gray), camera (dark green) and illuminator (light green). The inset shows a detail of the 3D-printed block (cyan) holding the response ports and allows appreciating the size and typical position of the dot’s pattern, relative to the other components of the rig.

**Figure 2.**
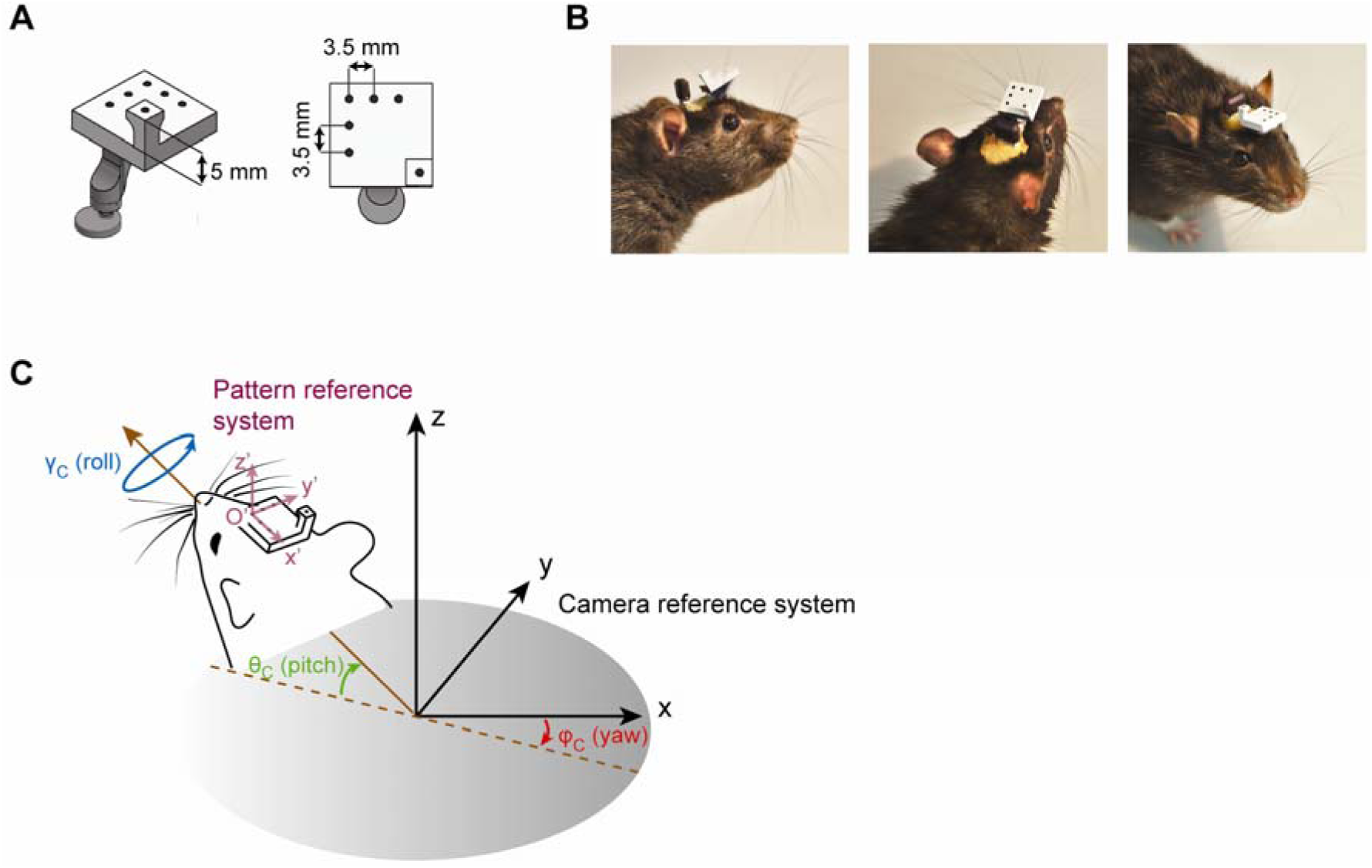
Illustration of the 3D pattern of dots used for head tracking and of the Euler angles that define its pose in the camera reference system. (**A**) A CAD rendering of the pattern of dots that, once mounted over the head of a rat, allows its position and pose to be tracked. Notice that, among the 6 dots, five are coplanar, while the sixth is placed over an elevated pillar. Also notice the arm that allows the pattern to be attached to a matching base surgically implanted over the skull of the rat. (**B**) Different views of a rat with the dots’ pattern mounted over his head. (**C**) Definition of the angles of rotation of the reference system centered on the dots’ pattern (*x*’, *y*’, *z*’; purple arrows) with respect to the camera reference system (*x, y, z*; black arrows). The three Euler angles – yaw, pitch and roll – are shown, respectively, by the red, green and blue arrows. *O*’ indicates the origin of the pattern reference system. The brown arrow indicates the direction where the rat’s nose is pointing and it is parallel to the head’s anteroposterior axis (i.e., to the *x*’ axis). The dashed brown line is the projection of the brown arrow over the (*x, y*) plane of the camera reference system.

It is more difficult (and only rarely it has been attempted) to achieve a complete estimate of the pose and location of the head in the three-dimensional (3D) space – i.e., to simultaneously track the three Cartesian coordinates of the head and the three Euler angles that define its orientation: yaw, pitch and *roll* (with the latter defined as the rotation about the head’s anteroposterior axis; see Figure 2C). Recently, two groups have successfully tracked in 3D the head of small, freely-moving mammals through videography, by relying either on a single camera imaging a custom tetrahedral arrangement of four LEDs with different colors (Finkelstein et al., 2015), or on multiple cameras (up to four) imaging custom 3D arrangements of up to six infrared (IR) LEDs (Sawinski et al., 2009; Wallace et al., 2013). Other groups have used instead inertial measurement units (IMUs), such as accelerometers and gyroscopes, mounted over the head of a rat, to record its angular displacement and velocity along the three Euler rotation axes (Pasquet et al., 2016; Kurnikova et al., 2017).

All these approaches provide accurate measurements of head position and pose in 3D. However, having been developed as ad hoc solutions for specific experimental settings, their design is not necessarily optimal for every application domain. For instance, most of these systems were conceived to track the head of small mammals roaming over an open-field arena, where the relatively large size of the custom LED- or IMU-based headset (extending several cm above and/or around the animal’s head) was not a issue in terms of encumbrance or obstruction. Moreover, these headsets need to be powered in order to operate. In general, this requires dedicated wires, which increase the stiffness of the bundle of cables connected to the headstage of the recording system, and prevent performing fully unplugged recordings using headstages equipped with wireless transmitters (Szuts et al., 2011; Pinnell et al., 2016).

In this study, we tried to overcome these limitations, by using a single, overhead camera to passively image a 3D-printed structure, painted with a pattern of black dots over a white background and mounted over the head of a rat. The small size of the pattern (1.35×1.35×1.5 cm) makes it ideal for perceptual studies, where a rodent performs a discrimination task inside a narrow operant box, often with its head inserted through an opening or confined within a funnel, as in the studies of rodent visual perception recently carried out by our group (Zoccolan et al., 2009; Tafazoli et al., 2012; Alemi-Neissi et al., 2013; Rosselli et al., 2015; Nikbakht et al., 2018; Djurdjevic et al., 2018) and other authors (Vermaercke and Op de Beeck, 2012; Horner et al., 2013; Mar et al., 2013; Kurylo et al., 2015; De Keyser et al., 2015; Bossens et al., 2016; Stirman et al., 2016; Kurylo et al., 2017; Yu et al., 2018). In what follows, beside describing in details the equipment and the algorithm upon which our method is based (Materials and Methods) and validate its accuracy and precision (first part of the Results and discussion), we provide a practical demonstration of the way our head tracker can help understanding: 1) how a rat samples the sensory stimuli during a visual or auditory discrimination task; and 2) how hippocampal neurons represent head position over extremely small spatial scales around the area where the animal delivers its perceptual decision and collects the reward.

## Results and discussion

Our head tracker was developed in the context of a two-alternative forced-choice (2AFC) perceptual discrimination experiment (Zoccolan, 2015; Zoccolan and Di Filippo, 2018), involving the visual and auditory modalities. A scheme of the experimental set up is shown in Figure 1A. The rats performed the discrimination task inside an operant box, similar to the one used in (Djurdjevic et al., 2018), and equipped with a monitor and two speakers for the presentation of the sensory stimuli. The task required each animal to insert the head through an opening in the wall facing the monitor and interact with an array of three response ports (i.e., three feeding needles, equipped with proximity sensors). Specifically, the rat had to lick the central needle to trigger the presentation of the stimulus. Afterward, he had to lick one of the lateral needles to report his perceptual choice and receive a liquid reward, in case of successful discrimination (see Materials and Methods).

Presentation of the visual and auditory stimuli, collection of the behavioral responses, delivery of reward, and real-time control of the flow of events during the experiment was achieved with the freeware, open source application MWorks (Figure 1A, solid black and cyan arrows), but any other software that allows programing psychophysical experiments would be suitable. Stimulus presentation was synchronized with the amplification/acquisition system for electrophysiological recordings, a Tucker-Davis Technologies (TDT) recording system (dotted black arrow) – i.e., at every trial, MWorks sent a code with the identity of the stimulus via UDP to the acquisition system.

The head tracker consisted of a far-red light source (red bulb) and an industrial monochromatic CMOS camera (green box), both placed above the operant box, with the camera feeding its video stream (green solid arrow) to a dedicated PC, equipped with the head tracking software. To synchronize the head tracker with the TDT acquisition system, we adopted a master-slave configuration, where the latter worked as a master, generating a square wave that triggered the camera image uptake (black dashed line). The camera, in turn, generated a unique identification code for every acquired frame, which was both saved by the PC running the head-tracking software (along with the Cartesian coordinates and pose of the head) and fed back to the TDT (green dashed arrow). In this way, the acquisition system simultaneously recorded the electric activity from the brain (red line), the identity and time of presentation of the stimuli displayed by MWorks, and the time and frame number of each image acquired by the overhead camera. Figure 1B shows a CAD drawing with a scaled representation of the key components of the rig and their relative position: the operant box (light gray), the monitor (dark gray), the block holding the feeding needles (cyan), the overhead camera (dark green) and the case of the light source (light green).

The key element of our head tracking systems is a 3D pattern of dots, mounted over the head of the animal and imaged by the overhead camera. The pattern is a small ((1.35×1.35×1.5 cm), light-weight (1.10 grams approximately), easy-to-place 3D printed structure, with 5 coplanar dots on a white background located over a plate, plus a sixth, elevated dot placed over a pillar (Figure 2A). At the beginning of each experimental session, the pattern is mounted on top of the head of the animal (Figure 2B), using an apposite magnet that connects it to a base that was previously surgically implanted (Materials and Methods). As described in detail in Materials and Methods, the head-tracking algorithm detects the 6 black dots in the frames of the video stream and computes the position and pose of the pattern in the 3D space of the camera reference system. More specifically, a *pattern reference system* (*x*’, *y*’, *z*’; purple arrows in Figure 2C) is defined, with the origin *O*’ placed over one of the dots, the *x*’ and *y*’s axes parallel to the edges of the plate, and the *z*’ axis perpendicular to it (i.e., parallel to the pillar). In the ideal case in which the pattern had been precisely aligned to the anatomical axes of the head at the time of the implant, *x*’ corresponds to the head’s anteroposterior axis, while *y*’ corresponds to the head’s interaural axis. The *camera reference system* (*x, y, z*; black arrows in Figure 2C) results instead from calibrating the camera using a standard procedure that consists in imaging a checkerboard pattern placed at various positions and orientations (see Materials and Methods). Once the camera is calibrated, the head-tracking algorithm provides: 1) the three Cartesian coordinates of *O*’ in the camera reference system; and 2) the rotation matrix *R* that defines the 3D rotation bringing the pattern reference system (*x*’, *y*’, *z*’) to be aligned with the camera reference system (*x, y, z*). *R* is defined as:

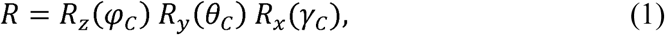

where *R*_*z*_, *R*_*y*_ and *R*_*x*_ are the elemental rotation matrixes that define intrinsic rotations by the Euler angles *φ*_*C*_ (yaw), *θ*_*C*_ (pitch), and *γ*_*C*_ (roll) about the axes of the camera reference system. More specifically:

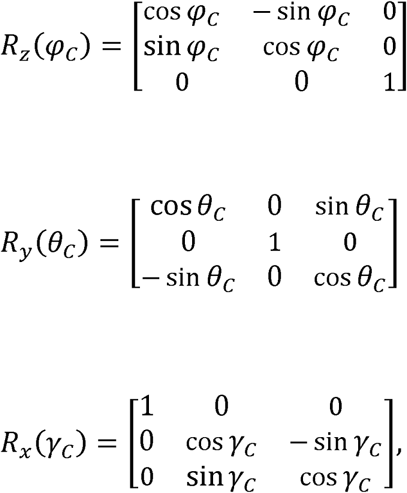

where, with reference to Figure 2C: *φ*_*C*_ is the angle between the projection of *x*’ onto the camera (*x, y*) plane and the camera *x* axis; *θ*_*C*_ is the angle between *x*’ and the (*x, y*) plane; and *γ*_*C*_ is the rotation angle of the pattern around *x*’.

It should be noticed that the three Euler angles *φ*_*C*_, *θ*_*C*_, and *γ*_*C*_, as well as the three Cartesian coordinates of *O*’, are not immediately applicable to know the pose and position of the head in the environment. First, they are relative to the camera reference system, while the experimenter needs to know them with respect to some meaningful environmental landmarks (e.g., the floor and walls of the arena or the operant box, where the animal is tested). Second, no matter how carefully the pattern is placed over the head of the rat, in general, the (*x*’, *y*’, *z*’) axes will not be perfectly aligned to the anatomical axes of the head – e.g., *x*’ and *y*’ will not be perfectly aligned to the head’s anteroposterior and interaural axes. However, expressing the pose and position of the pattern in camera coordinates allows measuring the nominal precision and accuracy that the head tracker can achieve, which is essential to validate the system. In the next section, we illustrate how we collected these validation measurements, while, in the following section, we explain how the actual position and pose of the head can be expressed in a convenient reference system, by collecting images of the checkerboard and dot patterns at, respectively, a reference position and pose.

### Validation of the head tracker: nominal precision and accuracy in the camera reference system

To measure the precision and accuracy of the head tracker (Figure 3), we used a custom combination of breadboards, linear stages, rotary stages and goniometers to hold the dots’ pattern in known 3D positions and poses. This allowed comparing the ground-true coordinates/angles of the pattern with the measurements returned by the head tracker. For these measurements, the camera was positioned in such a way to have its optical axis perpendicular to the floor of the testing area.

**Figure 3.**
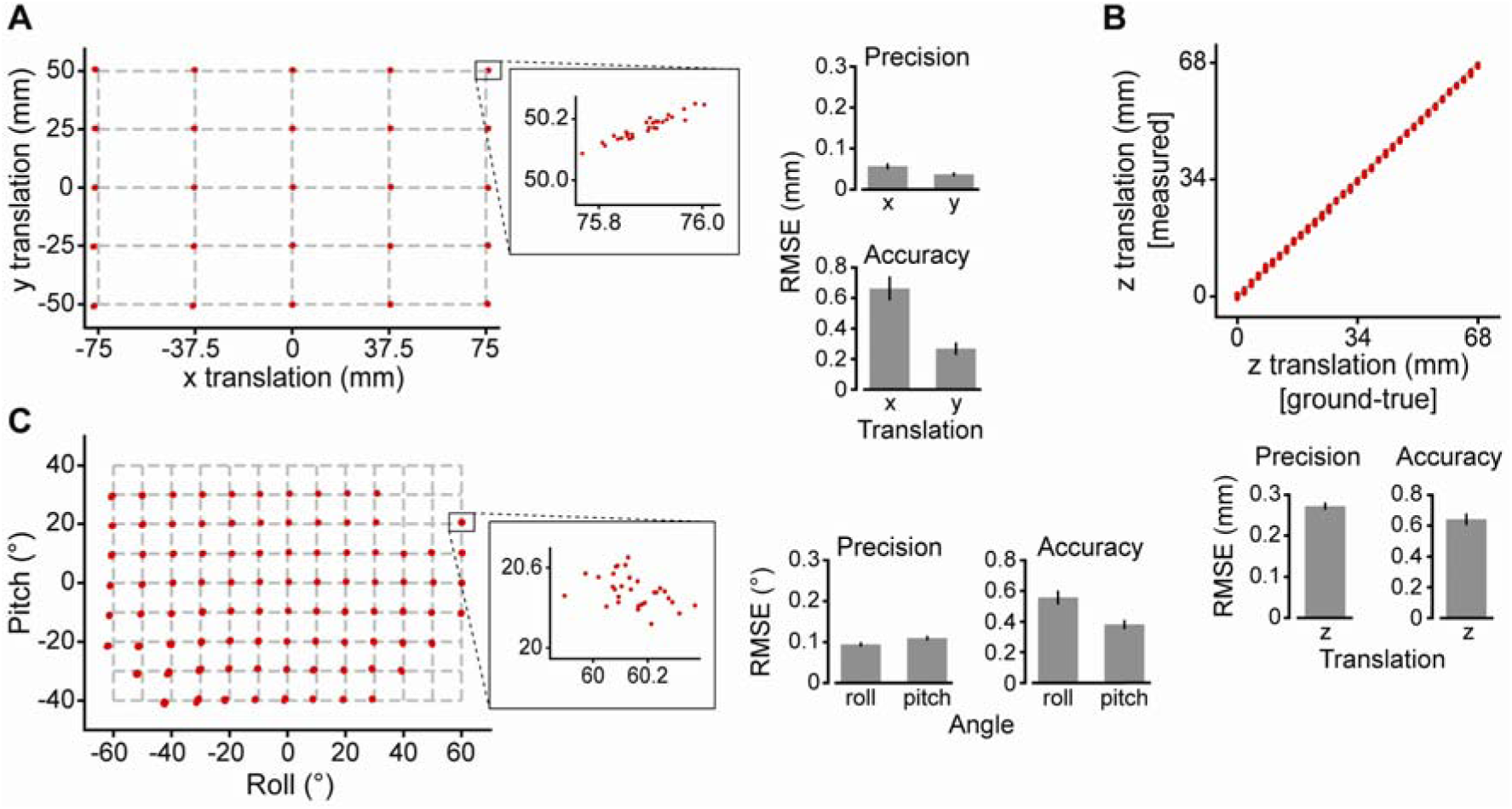
Nominal precision of the head-tracker in the camera reference system. (**A**) Left: validation measurements (red dots) obtained by placing the dots’ pattern on a grid of 5 × 5 ground-truth positions (grid’s intersections), over the floor of the testing area, using the breadboard shown in Figure 3-figure supplement 1A. Although 30 repeated head-tracker measurements were taken at each tested position, the spread of the measurements around the grid intersections is not appreciable, because of the very high precision of the head tracker. Only by zooming into the area of the gird intersection at the sub-millimetre scale, the spread of the red dots becomes visible (inset). Right: mean precision (top) and accuracy (bottom) of the head-tracker measurements over the 25 tested positions, as estimated by computing the RMSE relative either to the mean of each set of 30 repeated measurements (precision) or to the ground-truth positions (accuracy). Note that the sets of measured and ground-truth positions were aligned using Procrustes analysis (see main text). (**B**) Top: validation measurements (red dots) obtained by vertically displacing the dots’ pattern (relative to the floor of the testing area) of 34 consecutive increments, using the stereotax arm shown in Figure 3-figure supplement 1B. Again, given their high precision, the spread of the 30 repeated measurements taken at each ground-truth value is barely appreciable. Bottom: mean precision (left) and accuracy (right) of the head-tracker measurements over the 36 tested vertical displacements (RMSE computed as in **A**). (**C**) Left: validation measurements (red dots) obtained by setting the roll and pitch angles of the dots’ pattern to a combination of 13 × 9 ground-truth values (grid’s intersections), using the custom assembly shown in Figure 3-figure supplement 1C. The inset allows appreciating the spread of the 30 measurements taken at one of the tested angle combinations. Note that, for some extreme rotations of the pattern, no measurements could be taken (grid’s intersections without red dots), since the dots on the pattern were not visible. Right: mean precision (left) and accuracy (right) of the head-tracker measurements over the set of tested angle combinations (RMSE computed as in **A**). Note that the sets of measured and ground-truth angle combinations were optimally aligned following the procedure described in (Wallace et al., 2013) (see main text).

We first measured the ability of the system to track two-dimensional (2D) displacements of the pattern over the floor of the testing area. To this aim, the pattern was placed over a 3D printed breadboard with a grid of 5 × 5 holes (37.5 mm and 25 mm apart along, respectively, the horizontal and vertical dimensions; Figure 3-figure supplement 1A). For each of the 25 breadboard locations, we took a set of 30 repeated, head-tracker measurements of the origin of the pattern in the camera reference system (i.e., *O*’ in Figure 2C). Since the coordinates of the grid holes in such a reference system are not known a priori, a Procrustes analysis (Gower et al., 2004) was applied to find the optimal match between the set of 25×30 measures returned by the head tracker, and the known, physical positions of the holes of the breadboard. Briefly, the Procrustes analysis is a standard procedure to optimally align two shapes (or two sets of points, as in our application) by uniformly rotating, translating and scaling one shape (or one set of points) with respect to the other. In our analysis, since we compared physical measurements acquired with a calibrated camera, we did not apply the scale transformation (i.e., the scale factor fixed to 1). When applied to our set of 25×30 measures, the Procrustes analysis returned a very good match with the set of 25 ground-true positions of the grid (Figure 3A, left; red dots vs. grid intersections). As shown by the virtually absent spread of the dots at each intersection, the root mean square error (RMSE) of each set of 30 measurements, relative to their mean, was very low, yielding a mean *precision* (across positions) of 0.056 ± 0.007 mm along the x axis and 0.037 ± 0.005 mm along the y axis (Figure 3A, right; top bar plot). The RMSE of each set of 30 measurements, relative to the corresponding grid intersection, was also very low, yielding a mean *accuracy* (across positions) of 0.663 ± 0.079 mm along the x axis and 0.268 ± 0.041 mm along the y axis (Figure 3A, right; bottom bar plot). To verify the ability of the head-tracker to estimate the height of the pattern (i.e., its displacement along the z axis) when it was varied over a range of positions, the pattern was mounted on a stereotaxic arm through a 3D-printed custom joint (Figure 3-figure supplement 1B). The arm was positioned in such a way to be perpendicular to the floor of the testing area, thus allowing the pattern to be displaced vertically of 34 consecutive 2 mm increments. After each increment, the z coordinate of the pattern was estimated by the head tracker in 30 repeated measurements, which were then compared to the total physical displacement of the stereotaxic arm up to that point (Figure 3B, top). The resulting estimates were again very precise and accurate (Figure 3B, bottom), with RSME values that were close to those obtained previously for the x and y displacements (compare the z bars of Figure 3B to the x and y bars of Figure 3A).

For the validation of the angular measurements, we built a custom assembly, made of a rotary stage mounted on a stereotaxic arm, that allowed rotating the pattern about two orthogonal axes, corresponding to roll and pitch (Figure 3-figure supplement 1C). Rotations about each axis were made in 10° steps, spanning from −60° to 60° roll angles and from −40° to 40° pitch angles, while the yaw was kept fix at 0°. Again, 30 repeated head-tracker measurements were collected at each know combination of angles over the resulting 13 × 9 grid. As for the case of the 2D displacements, the angles returned by the head tracker were not immediately comparable with the nominal rotations on the rotary stages, because the two sets of angles are measured with respect to two different reference systems – i.e., the camera reference system (as defined in Figure 2C), and the stage reference system (as defined by the orientation of the pattern in the physical environment, when the rotations of the stages are set to zero). Therefore, in order to compare the nominal rotations of the pattern on the stages 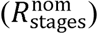 to their estimates provided by the head-tracker 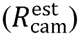, we first had to express the former in the camera reference system 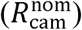. To this aim, we followed the same approach of (Wallace et al., 2013) and we computed 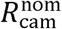 as:

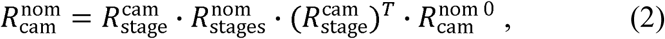

where 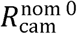 is the pose of the pattern (in the camera reference system) when all the rotations of the stages are nominally set to zero (i.e., reference rotation), and 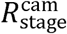 is the matrix mapping the stage reference system into the camera reference system [note that each matrix in (2) is in the form shown in (1)]. The matrixes 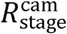 and 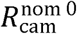 are unknown that can be estimated by finding the optimal match between the nominal and estimated rotations in camera coordinates, i.e., between 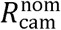, as defined in (2), and 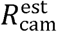. Following (Wallace et al., 2013), we defined the rotation difference matrix 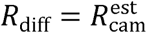·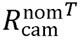, from which we computed the error of a head-tracker angle estimate as the total rotation in *R*_diff_, i.e., as:

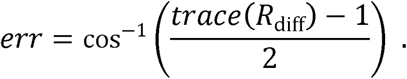

By minimizing the sum of *err*^2^ over all tested rotations of the stages (using Matlab *fminsearch* function), we obtained a very close match between estimated and nominal stage rotations. This is illustrated in Figure 3C (left), where the red dots are the head-tracker estimates and the grid intersections are the nominal rotations. As for the case of the Cartesian displacements, also for the pitch and roll angles, the head tracker returned very precise (roll: RSME = 0.095 ± 0.005°; pitch: RSME = 0.109 ± 0.006°) and accurate (roll: RSME = 0.557 ± 0.045°; pitch: RSME = 0.381± 0.030°) measurements (Figure 3C, right).

### Operation of the head tracker: measuring displacements and rotations relative to reference poses in the physical environment

While the validation procedure described in the previous section provides an estimate of the nominal precision and accuracy of the head tracker, measuring Cartesian coordinates and Euler angles in the camera reference system is impractical. To refer the head tracker measurements to a more convenient reference system in the physical environment, we 3D printed a custom adapter to precisely place the checkerboard pattern used for camera calibration over the block holding the feeding needles, in such a way to be parallel to the floor of the operant box, with the vertex of the top-right black square vertically aligned with the central needle (see Figure 4-figure supplement 1A). We then acquired an image of this *reference checkerboard*, which served to establish a new reference system (*x*”, *y*”, *z*”), where the *x*” and *y*” axes are parallel to the edges at the base (the floor) of the operant box, while *z*” is perpendicular to the floor and passes through the central feeding needle. The position measurements returned by the head tracker can be expressed as (*x*”, *y*”, *z*”) coordinates by applying the rotation matrix 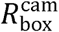 and the translation vector 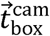 that map the camera reference system into this new operant box reference system. 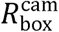 is in the form shown in (1), but with the angles referring to the rotation of the reference checkerboard with respect to the camera reference system; 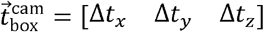, where Δ*t* is the distance between the origins of the two reference systems along each axis.

**Figure 4.**
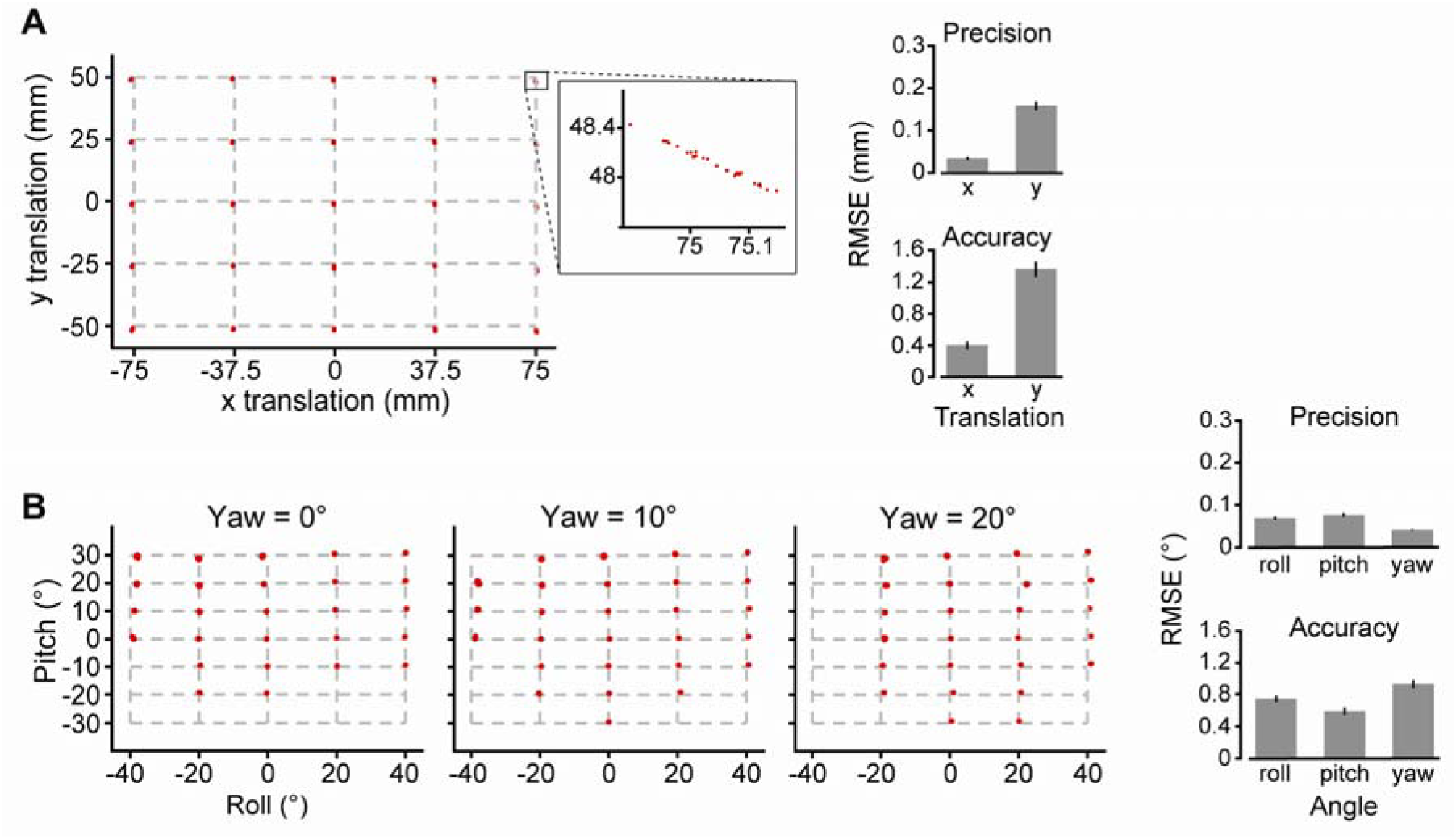
Validation of the head-tracker in the reference system of the operant box. (**A**) Left: validation measurements (red dots) obtained by placing the dots’ pattern on a grid of 5 × 5 ground-truth positions (grid’s intersections; same as in Figure 3A), over the floor of the operant box. Each measurement is relative to a *box reference system*, with the *x* and *y* axes parallel to the floor of the operant box, and the z axis passing through the central response port (see Figure 1A). Converting the measurements from the camera to the box reference system was possible by acquiring an image of a reference checkerboard, as described in the main text and shown in Figure 4-figure supplement 1A. The inset allows appreciating the spread of the 30 measurements taken at one of the tested positions. Right: mean precision (top) and accuracy (bottom) of the head-tracker measurements over the 25 tested positions (RMSE computed as in Figure 3A). (**B**) Left: validation measurements (red dots) obtained by setting the yaw, pitch and roll angles of the dots’ pattern to a combination of 3 × 7 × 5 ground-truth values (grid’s intersections), using the custom assembly shown in Figure 4-figure supplement 1B. Each measurement is relative to a *pose zero reference system*, obtained by acquiring an image of the dots’ pattern with all the angles on custom assembly set to zero. Note that, given their high precision, the spread of the 30 repeated measurements taken at each ground-truth value is barely appreciable. Right: mean precision (left) and accuracy (right) of the head-tracker measurements over the set of tested angle combinations (RMSE computed as in Figure 3A).

We tested the ability of the head tracker to correctly recover the Cartesian coordinates of the dots’ pattern relative to the box reference system, by placing the pattern over the 5×5 grid shown in Figure 3-figure supplement 1A and collecting 30 repeated, head-tracker measurements at each location. To verify the functioning of the head tracker under the same settings used for the behavioral and neurophysiological experiments (see next sections), the camera was not centered above the operant box, with the optical axis perpendicular to the floor, as previously done for the validation shown in Figure 3. Otherwise, during a neuronal recording session, the cable that connects the headstage protruding from the rat head to the preamplifier would partially occlude the camera’s field of view. Hence, the need to place the camera in front of the rat, above the stimulus display, oriented with an angle of approximately 50° relative to the floor (see Figure 1B). This same positioning was used here to collect the set of validation measurements over the 5×5 grid. As shown in Figure 4A, the match between estimated and nominal horizontal (*x*”) and vertical (*y*”) coordinates of the pattern was very good (compare the red dots to the grid intersection), with a barely appreciable dispersion of the 30 measurements around each nominal position value. This resulted in a good overall precision (x: RMSE = 0.034 ± 0.004; y: 0.158 ± 0.010) and accuracy (x: RMSE = 0.403 ± 0.050; y: 1.36 ± 0.093) and of the x” and y” measurements.

The Euler angles defining the pose of the head in the 3D space could also be measured relative to the operant box reference system (*x*”, *y*”, *z*”). However, this would yield an estimate of the rotation of the dots’ pattern, rather than of the rat head, in the physical environment. In fact, no matter how carefully the support holding the pattern is implanted at the time of the surgery, it is unlikely for the pattern to be perfectly aligned to the anatomical axes of the head, once put in place. In general, the *x*’ and *y*’ axes of the pattern reference system (see Figure 2B) will be slightly titled, with respect to the head’s anteroposterior and interaural axes. Therefore, it is more convenient to acquire an image of the pattern when the rat is placed in the stereotax, with its head parallel to the floor of the operant box, and then use such a *pose zero* of the pattern as a reference for the angular measurements returned by the head tracker. This can be achieved by defining a pose zero reference system 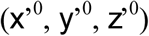 and a rotation matrix 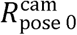 mapping the camera reference system into this new coordinate system [the matrix is in the form shown in (1), but with the angles referring to the rotation of the pose zero of the pattern with respect to the camera reference system].

To test the ability of the head tracker to correctly recover the Euler angles of the dots’ pattern relative to the pose zero reference system, we mounted the pattern over a custom assembly made of two 3D printed goniometers, each allowing rotations over a span of 70° (from −35° to +35°), and a rotary stage, enabling 360° rotations (Figure 4-figure supplement 1B). This allowed setting the pose of the pattern over a grid of known 3×7×5 combinations of yaw, pitch and roll angles. As illustrated in Figure 4B (left), we found a good match between the nominal and estimated pattern rotations (note that not all 3×7×5 angle combinations were actually tested, since the dots on the pattern were not detectable at some extreme rotations). When mediated across all tested angle combinations, the resulting precision and accuracy (Figure 4B, right) were very similar to the nominal ones shown in Figure 3C (precision: roll 0.068 °± 0.003°, pitch 0.076° ± 0.004°, yaw 0.0409° ± 0.001°; accuracy: roll 0.746° ± 0.036, pitch 0.598° ± 0.048°, yaw 0.929° ± 0.052°).

### Head movements of a rat performing a two-alternative forced choice task in the visual and auditory modalities

To illustrate the application of our head-tracker *in-vivo*, we implanted a rat with an electrode array targeting hippocampus (see Materials and Methods and next section for details). The implant also included a base with a magnet that allowed attaching the dots’ pattern during the behavioral/recording sessions (see Figure 2A-B). As previously explained, the animal had to interact with an array of three response ports, each equipped with a feeding needle and a proximity sensor (Figure 1). Licking the central port triggered the presentation of either a visual object (displayed on the screen placed in front of the rat) or a sound (delivered through the speakers located on the side of the monitor). Two different visual objects could be presented to the animal [same as in (Zoccolan et al., 2009; Alemi-Neissi et al., 2013a)] – Object 1 required the rat to approach and lick the left response port in order to obtain liquid reward, while Object 2 required him to lick the right response port (Figure 5A). The same applied to the two sounds, one associated to the left and the other to right response port.

**Figure 5.**
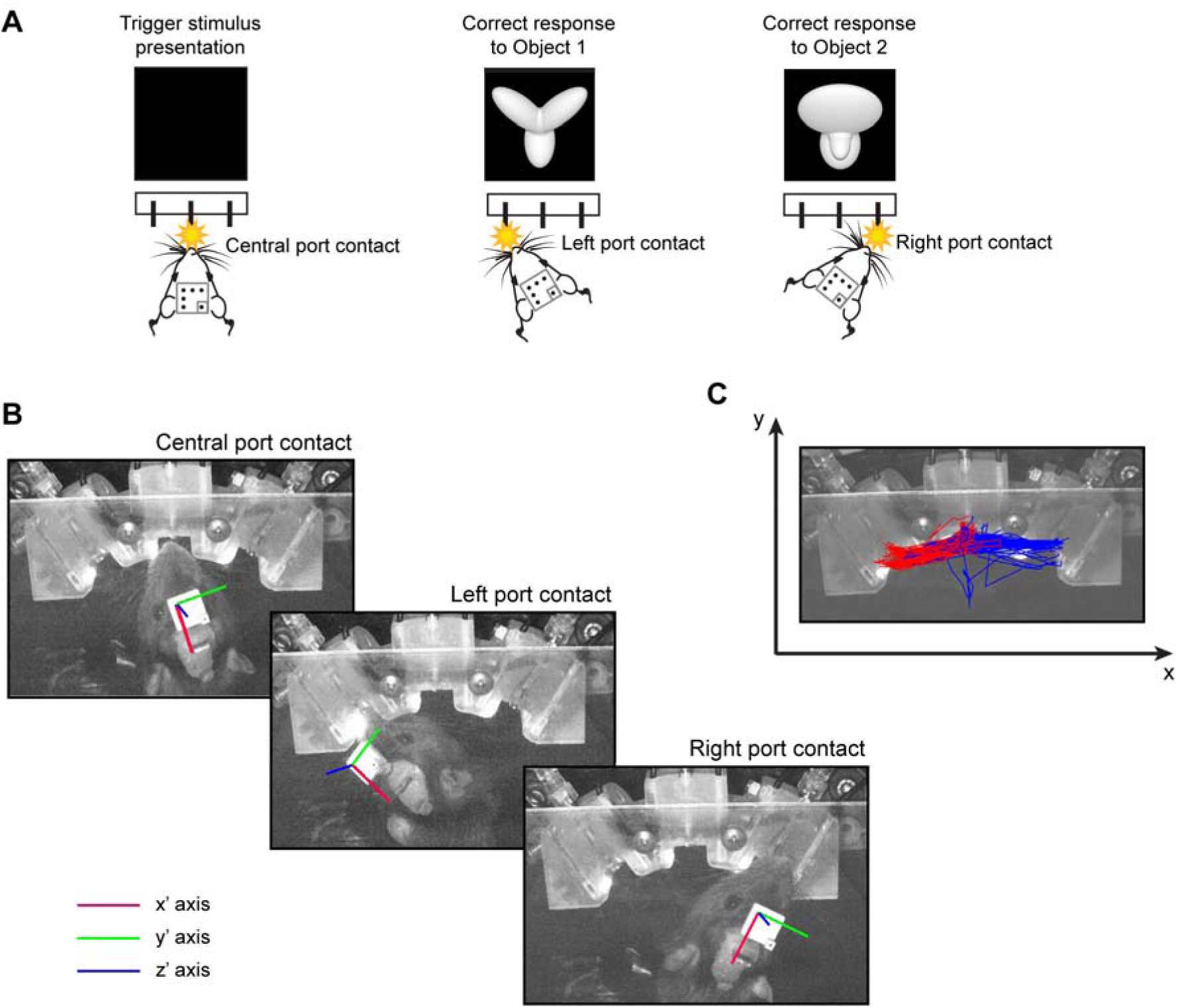
Head tracking of a rat engaged in a perceptual discrimination task. (**A**) Illustration of the visual discrimination task. The rat learned to lick the central response port to trigger the presentation of either Object 1 or Object 2 on the stimulus display placed in front of the operant box (see Figure 1A). Presentation of Object 1 required the rat to approach and lick the response port on the left in order to correctly report its identity, while presentation of Object 2 required the animal to lick the response port on the right. (**B**) Example snapshots captured and processed by the head tracker at three representative times – i.e., when the rat licked the central, the left and the right response ports. The colored lines are the x’ (red), y’ (green) and z’ (blue) axes of the reference system centered on the dots’ pattern (se Figure 2C), as inferred in real time by the head-tracker (see also supplementary Figure 5-video supplement 1). (**C**) The trajectories followed by the nose of the rat in consecutive trials during the execution of the task are superimposed to a snapshot of the block holding the response ports, as imaged by the head tracker. The red and blue traces refer to trials in which the animal chose, respectively, the left and right response port. The trajectories are plotted in the Cartesian plane corresponding to the floor of the operant box, where the *x* and *y* axes (black arrows) are, respectively, parallel and orthogonal to the stimulus display (see also Figure 1A).

Figure 5B shows example images captured and processed by the head-tracker in three representative epochs during the execution of the task (the colored lines are the *x*’, *y*’ and *z*’ axes that define the pose of the pattern, as inferred in real time by the head-tracker; see supplementary Figure 5-video supplement 1). By tracking the position of the pattern in the 2D plane of the floor of the operant box, it was possible to plot the trajectory of the rat’s nose in each individual trial of the behavioral test (Figure 5C, red vs. blue lines, referring to trials in which the animal chose, respectively, the left and right response port). The position of the nose was monitored, because it better reflects the interaction of the animal with the response sensors, as compared to the position of the pattern (the latter can be converted into the nose position by carefully measuring the distance between the origin of the pattern and the tip of the nose at the time of the surgery). With such a precise knowledge of the position and pose of the nose/head in space and time, we could address questions concerning the perceptual, motor and cognitive processes deployed by the rat during the visual and auditory discriminations, well beyond the knowledge that can be gained by merely monitoring the response times collected by the proximity sensors.

For example, in Figure 6A (left), we have reported the x position of the rat’s nose as a function of time in all the behavioral trials collected over 14 consecutive test sessions (total of 3,413 trials). The traces are aligned to the time in which the stimulus was presented (i.e., 300 ms after the animal had licked the central port). From these traces, we computed the *reaction time* (RcT) of the rat in each trial, defined as the time, relative to the stimulus onset, in which the animal left the central sensor to start a motor response, eventually bringing its nose to reach either the left (red lines) or right (blue line) response port. As it can be appreciated by looking at the spread of both sets of traces, RcT was highly variable across trials, ranging from 300 ms to 1166 ms (extreme values not considering outliers), with a median around 530 ms (Figure 6A, right). By contrast, a much smaller variability was observed for the *ballistic response time* (BRsT), i.e., the time taken by the rat to reach the left or right response port, relative to the last time he visited (i.e., passed by) the central port (Figure 6B, left), with BRsT ranging between 66 ms and 566 ms (Figure 6B, right: upper box plot; median ~300 ms). This suggests that much of the variability in the *total response time* (ToRsT; i.e., the time taken to reach the left or right response port, relative to the stimulus onset; Figure 6B, right: bottom box plot) has to be attributed to the perceptual/decisional process required for the correct identification of the stimulus. However, a close inspection of the response trajectories of Figure 6B (aligned to the time in which the ballistic motor response was initiated) shows that the rat, after leaving the central port in response to the stimulus, often did not point directly to the port that he would eventually choose (referred to as the *selected response port* in what follows). In many trials, the final ballistic response was preceded by earlier movements of the head, either towards the selected response port or the opposite one (referred to as the *opposite response port* in what follows).

**Figure 6.**
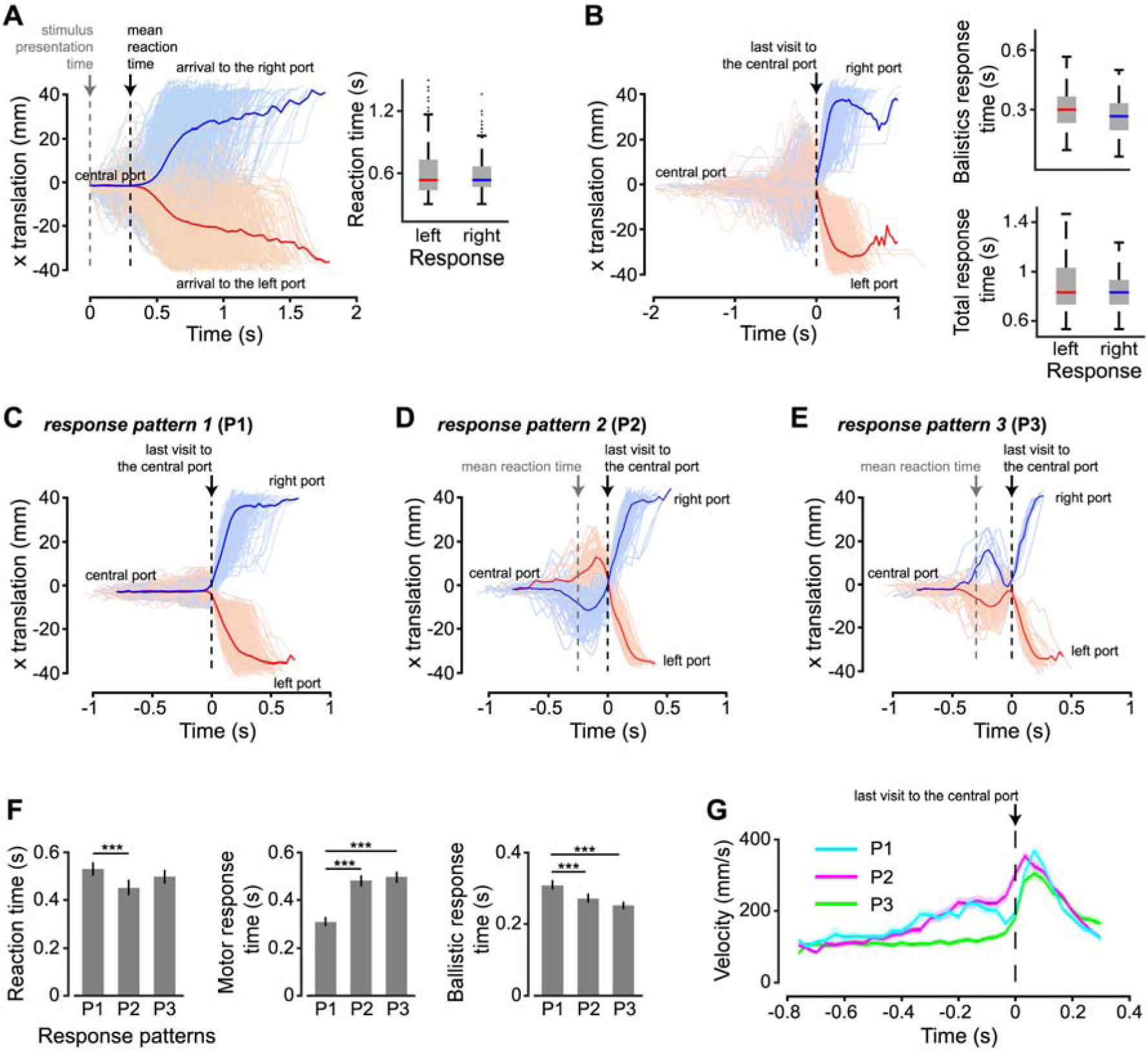
Statistical characterization of the head’s displacements performed by the rat during the perceptual discrimination task. (**A**) Left: position of the rat’s nose as a function of time, along the *x* axis of the Cartesian plane corresponding to the floor of the operant box (i.e., *x* axis in Figure 5C). The traces, recorded in 3,413 trials over the course of 14 consecutive sessions, are aligned to the time in which the stimulus was presented (gray dashed line). The black dashed line indicates the mean reaction time (as defined in the main text). The red and blue colors indicate trials in which the rat chose, respectively, the left and right response port. Right: box plot showing the distributions of reaction times for the two classes of left and right responses. (**B**) Left: same trajectories as in **A**, but aligned to the last time the rat visited the central port (dashed line). Right: box plots showing the distributions of ballistic response times (top) and total response times (bottom) for the two classes of left and right responses (see main text for definitions). (**C**) Subset of the response patterns (referred to as *P1*) shown in **A** and **B**, in which the rat, after leaving the central port, made a direct ballistic movement to the selected response port (traces’ alignment as in **B**). (**D**) Subset of the response patterns (referred to as *P2*) shown in **A** and **B**, in which the rat, before making a ballistic movement towards the selected response port, made an initial movement towards the opposite port (traces’ alignment as in **B**). (**E**) Subset of the response patterns (referred to as *P3*) shown in **A** and **B**, in which the rat made an initial movement towards the selected response port, then moved back to the central port, and finally made a ballistic moment to reach the selected port (traces’ alignment as in **B**). (**F**) Comparisons among the mean reaction times (left), among the mean motor response times (middle; see main text for a definition) and among the mean ballistic response times (right) that were measured for the three types of trajectories (i.e., P1, P2 and P3) shown, respectively, in **C**, **D** and **E** (*** *p* < 0.001; two-tailed, unpaired t-test). Error bars are SE of the mean. (**G**) Velocity of the rat’s nose as a function of time for the three types of trajectories P1, P2 and P3.

To better investigate rat response patterns, we separated the recorded trajectories into three response patterns. In the first response pattern (P1), the rat, after leaving the central port, made a direct ballistic movement to the selected response port (either left or right; Figure 6C). In the second response pattern (P2), the rat made an initial movement towards the opposite response port, before correcting himself and making a ballistic movement towards the selected response port (Figure 6D). In the third response pattern (P3), the rat made an initial movement towards the selected response port, but then moved back to the central port, before approaching again, with a ballistic moment, the selected port (Figure 6E). Interestingly, RcT was significantly smaller in P2 than in P1 (*p* < 0.001; two-tailed, unpaired t-test; Figure 6F, leftmost bar plot), suggesting that the trials in which the animal reversed his initial decision were those in which he made an “impulsive” choice that he eventually corrected. As expected, the *motor response time* (MoRsT; i.e., the time taken by the animal to reach the selected response port, after leaving, for the first time, the central port) was substantially lower in P1, as compared to P2 and P3, given the indirect trajectories that the latter trial types implied (*p* < 0.001; two-tailed, unpaired t-test; Figure 6F, central bar plot). By contrast, BaRsT was slightly, but significantly higher in P1 than in P2 and P3, indicating that ballistic movements were faster, when they followed a previously aborted choice (*p* < 0.001; two-tailed, unpaired t-test; Figure 6F, rightmost bar plot).

To better understand this phenomenon, we plotted the average velocity of the rat’s nose as a function of time for the three response patterns (Figure 6G; curves are aligned to the onset of the ballistic motor response; dashed line). As expected, in P1 (green curve), the velocity was close to zero till the time in which the ballistic response was initiated. By contrast, in P2 (magenta curve), the velocity was already high at the onset of the ballistic movement, because the animal’s nose passed by the central sensor when sweeping from the opposite response port to the selected one (see Figure 6D). This momentum of the rat’s head was thus at the origin of the faster ballistic responses in P2, as compared to P1. In the case of P3 (cyan curve), there was no appreciable difference of velocity with P1 at the onset of the ballistic movements. This is expected, given that the rat moved twice in the direction of the selected port and, in-between the two actions, he stopped at the central sensor (see Figure 6E). However, following the onset of the ballistic movement, the rat reached a larger peak velocity in P3 than in P1, which explains the shorter time needed to complete the ballistic response in the former trials’ type. This may possibly indicate a larger confidence of the rat in his final choice, following an earlier, identical response choice that was later aborted.

We also used the head tracker to inquire whether the rat deployed different response/motor patterns depending on the sensory modality of the stimuli he had to discriminate. Rat performance was higher in the sound discrimination task, as compared to the visual object discrimination task (*p* < 0.001; two-tailed, unpaired t-test; Figure 7A). Consistently with this observation, the fraction of trials in which the animal aborted an initial perceptual choice (i.e., response patterns P2 and P3), as opposed to make a direct response to the selected port (i.e., response pattern P1), was significantly larger in visual than in auditory trials (*p* < 0.01, χ^2^ test for homogeneity; Figure 7B). This means that the animal was less certain about his initial decision in the visual trials, displaying a tendency to correct more often such decision, as compared to the auditory trials. Interestingly, the lower perceptual discriminability of the visual stimuli did not translate into a general tendency of reaction times to be longer in visual than auditory trials. When the animal aimed directly to the selected port (P1; the vast majority of trials, as it can be appreciated in Figure 7B), no significant difference in RcT was observed between visual and auditory trials (*p* > 0.05; two-tailed, unpaired t-test; Figure 7C, top-left bar plot). By contrast, in trials in which the rat corrected his initial decision (P2), RcT was significantly longer in visual than auditory trials (*p* < 0.01; two-tailed, unpaired t-test; Figure 7C, middle-left bar plot), but the opposite trend was found in trials in which the animal swung back and forth to the selected response port (P3; *p* < 0.05; two-tailed, unpaired t-test; Figure 7C, bottom-left bar plot). We found instead a general tendency of the rat to make faster ballistic responses in visual than auditory trials, with this trend being significant in P1 and P2 (*p* < 0.001; two-tailed, unpaired t-test; Figure 7D, top-right and middle-right bar plots). It is hard to interpret these findings, which could indicate a larger impulsivity of the animal in the visual discrimination, but, possibly, also a larger confidence in his decision. Addressing more in depth this issue is obviously beyond the scope of our study, since it would require measuring the relevant metrics (e.g., RcT and BaRsT) over a cohort of animals, while our goal here was simply to provide an in-vivo demonstration of the working principle of our head-tracker and suggest possible ways of using it to investigate the perceptual, decision and motor processes involved in a perceptual discrimination task.

**Figure 7.**
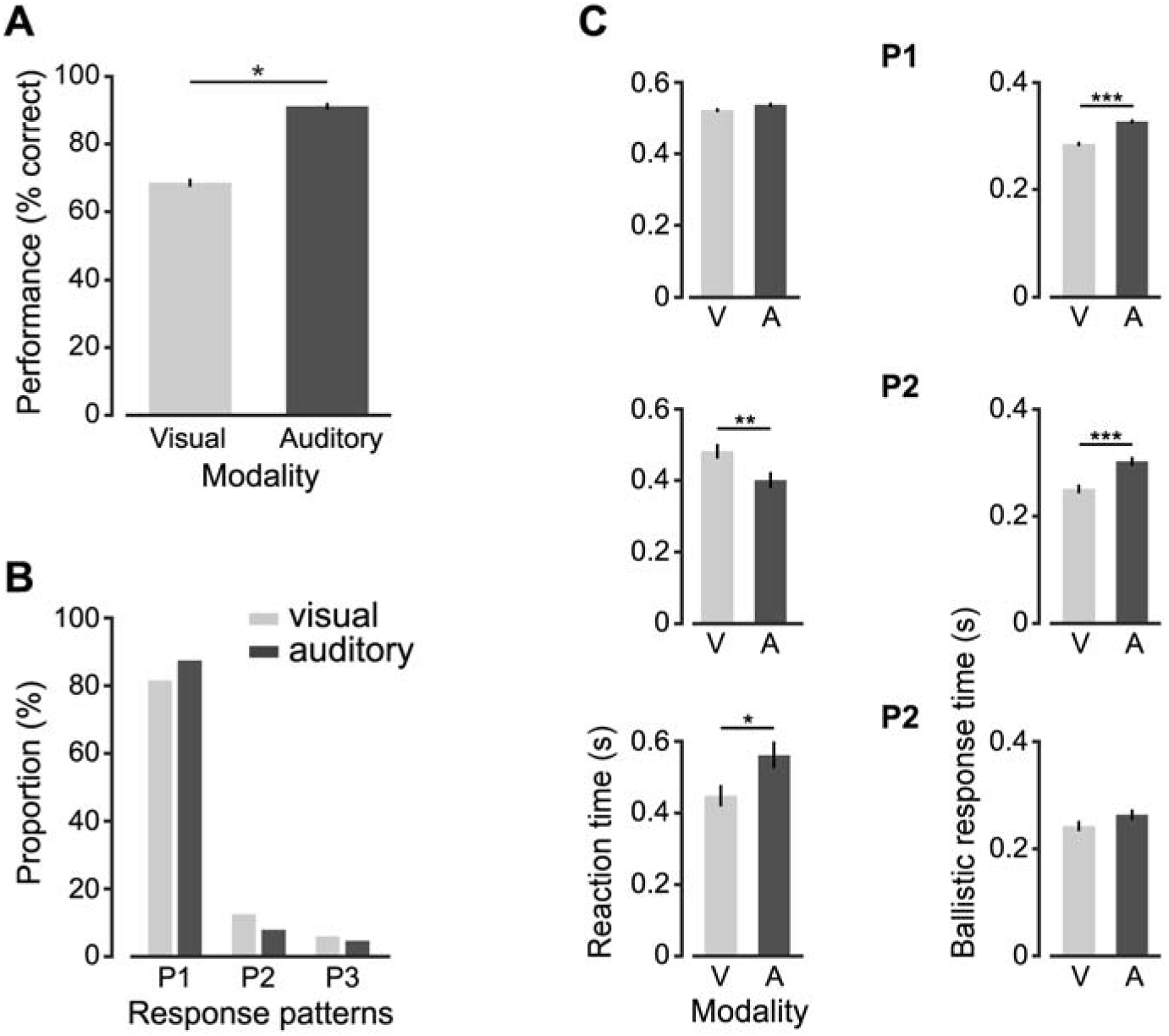
Statistical comparison of the head’s displacements performed by the rat in visual and auditory trials. (**A**) Comparison between the performances attained by the rat in the visual (light gray) and auditory (dark gray) discrimination tasks (*** *p* < 0.001; two-tailed, unpaired t-test). (**B**) Proportion of response patterns of type P1, P2 and P3 (see Figure 6C-E) observed across the visual (light gray) and auditory (dark gray) trials. The two distributions were significantly different (*p* < 0.01, X^2^ test). (**C**) Comparisons between the mean reaction times (left) and between the mean ballistic response times (right) that were measured in visual (light gray) and auditory (dark gray) trials for each type of trajectories (i.e., P1, P2 and P3). * *p* < 0.05, ** *p* < 0.01, *** *p* < 0.001; two-tailed, unpaired t-test.

As a further example of the kind of behavioral information that can be extracted using the head-tracker, we analyzed the pose of the rat’s head during the execution of the task (Figure 8). As shown in Figure 8B, at the time the rat triggered the stimulus presentation (time 0), his head was, on average across trials (thick curves), parallel to the floor of the operant box (i.e., with pitch 0° and roll close to 0°) and facing frontally the stimulus display (i.e., with yaw close to 0°). However, at the level of single of trials (thin curves), the pose of the head was quite variable, with approximately a ±30° excursion in the pitch, and a ±15°/20° excursion in the roll and yaw. This can be also appreciated by looking at the polar plots of Figure 8C (central column), which report the average pitch, roll and yaw angles that were measured over the 10 frames (~300 ms) following the activation of each response port (thin lines: individual trials; thick lines: trials’ average). Being aware of such variability is especially important in behavioral and neurophysiological studies of rodent visual perception (Zoccolan, 2015). In fact, a variable pose of the head at the time of stimulus presentation implies that the animal viewed the stimuli under quite different angles across repeated behavioral trials. This, in turns, means that the rat had to deal with a level of variation in the appearance of the stimuli on his retina that was larger than that imposed, by the design, by the experimenter. In behavioral experiments where the invariance of rat visual perception is under investigation, this is not an issue, because, as observed in (Alemi-Neissi et al., 2013), it can lead at most to an underestimation (not to an overestimation) of rat invariant shape-processing abilities. However, in studies were a tight control over the retinal image of the visual stimuli is required, the trial-by-trial variability reported in Figure 8B-C indicates that the use of a head-tracker is necessary to measure, and possibly compensate, the change of viewing angle occurring across repeated stimulus presentations. This applies, for instance, to neurophysiological studies of visual representations in unrestrained (i.e., not head-fixed) rodents, especially when localized stimuli (e.g., visual objects) are used to probe low- and middle-level visual areas. For instance, head-tracking, ideally also paired with eye-tracking, would be necessary to investigate putative ventral stream areas in unrestrained rats, as recently done in anesthetized (Tafazoli et al., 2017; Matteucci et al., 2019) or awake, but head-fixed animals (Vinken et al., 2014, 2016, 2017; Kaliukhovich and Op de Beeck, 2018). By contrast, the issue of pose variability afflicts less neurophysiological studies targeting higher-order association or decision areas, especially when using large (ideally full-field) periodic visual patterns (e.g., gratings) (Nikbakht et al., 2018).

**Figure 8.**
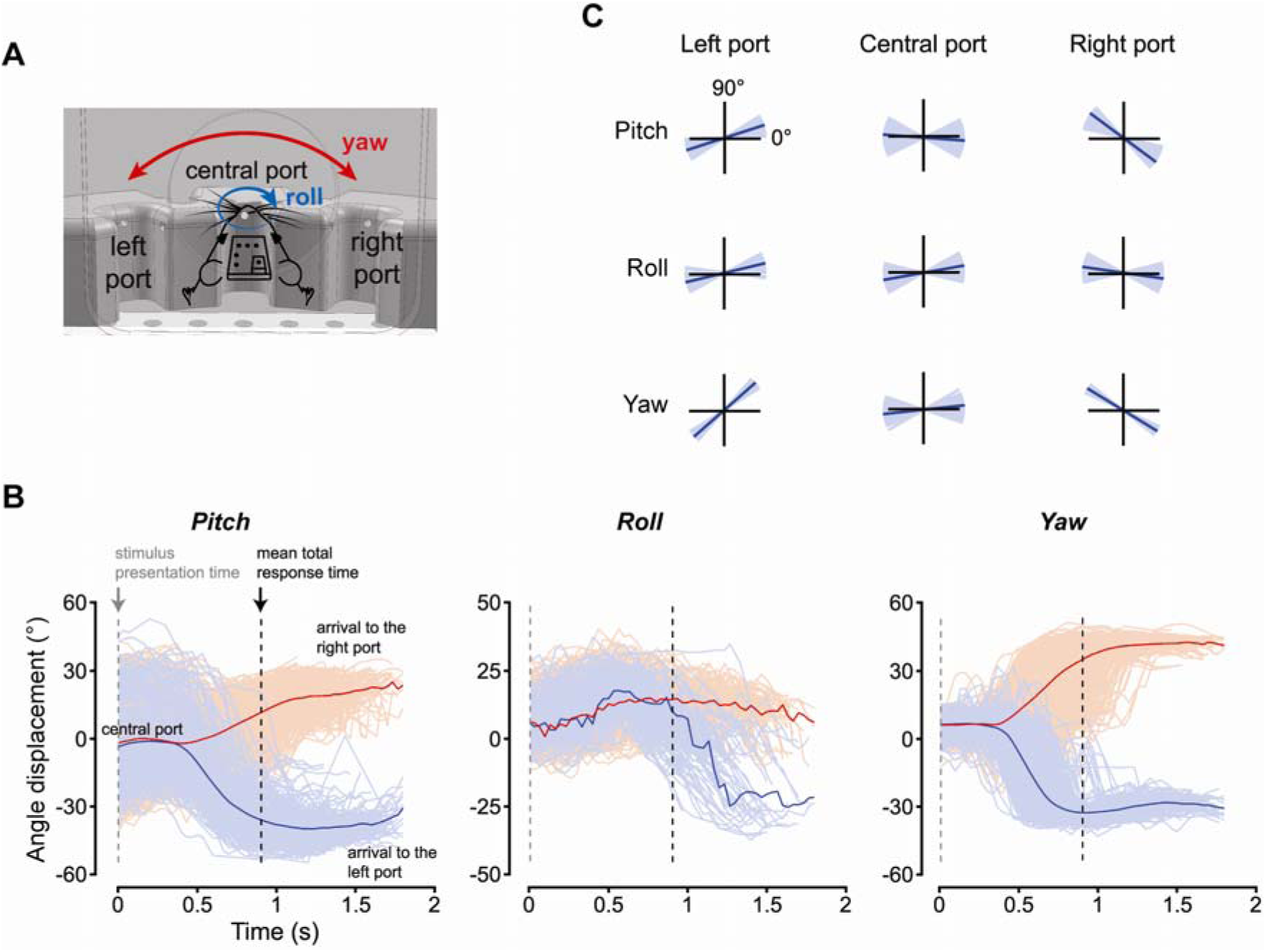
Statistical characterization of the head’s rotations performed by the rat during the perceptual discrimination task. (**A**) Illustration of the yaw and roll rotations that the rat’s head can perform, relative to its reference pose (see main text). (**B**) Time course of the pitch (left), roll (middle) and yaw (right) angles during repeated trials of the perceptual discrimination task. The red and blue colors indicate trials in which the rat chose, respectively, the left and right response port. Traces are aligned to the time in which the stimulus was presented (dashed gray line). The mean total response time (i.e., the mean time of arrival to the selected response port) is shown by the dashed black line. (**C**) The pitch (top), roll (middle) and yaw (bottom) angles measured by the head tracker in individual trials (thin blue lines) and, on average, across all trials (thick black lines), at the times the rat reached the left, central and right response ports.

Monitoring the pose of the head during the behavioral trials also revealed that the rat approached the lateral responses ports with his head at port-specific angles (compare red vs. blue curves in Figure 8B, and left vs. right columns in Figure 8C). Unsurprisingly, the yaw angle was the opposite for left (about 40°) and right (about −35°) responses, since the animal had to rotate his head towards opposite direction to reach the lateral response ports (see Figure 8A, red arrows). It was less obvious to observe opposite rotations also about the roll and pitch axes, which indicate that the rat bent his head in port-specific ways to reach each response port and lick from the corresponding feeding needle. Again, this information is relevant for behavioral studies of rodent visual perception, where the stimulus is often left on the screen after the animal makes a perceptual choice and during the time he retrieves the liquid reward (Zoccolan et al., 2009; Alemi-Neissi et al., 2013; Rosselli et al., 2015; Djurdjevic et al., 2018). This implies that the animal experiences each stimulus from a port-specific (and therefore stimulus-specific) viewing angle for a few seconds after the choice. As pointed out in (Djurdjevic et al., 2018), this can explain why rats learn to select specific stimulus features to process a given visual object.

### Simultaneous head tracking and neuronal recordings during a two-alternative forced choice discrimination task

To illustrate how our head tracker can be combined with the recording of neuronal signals, we monitored the head movements of the rat during the execution of the visual/auditory discrimination task, while recording the activity of hippocampal neurons in CA1. Given that, in rodents, hippocampal neurons often code the position of the animal in the environment (Moser et al., 2008, 2015), we first built a map of the places visited by the rat while performing the task (Figure 9A, top). It should be noticed that, differently from typical hippocampal studies, where the rodent is allowed to freely move inside an arena, the body of the rat in our experiment was at rest, while his head, after entering through the viewing hole, made small sweeps among the three response ports. Therefore, the map of visited positions shown in Figure 9A refers to the positions of the rat’s nose, rather than to his body. Thus, by binning the area around the response ports, we obtained a density map of visited locations by the nose (Figure 9A, bottom). Not surprisingly, this map revealed that the rat spent most of the time with his nose very close to one of the response ports (red spots). Figure 9B shows instead the average firing rate of an example, well-isolated hippocampal neuron (whose waveforms and inter-spike interval distribution are shown in Figure 9C) as a function of the position of the rat’s nose (only locations that were visited more than 10 times were considered). The resulting spatial map displayed a marked tendency of the neuron to preferentially fire when the rat approached the right response port. In other words, this neuron showed the sharp spatial tuning that is typical of hippocampal place cells, but over a much smaller spatial extent (i.e., over the span of his head movements around the response ports) than typically observed in freely moving rodents.

**Figure 9.**
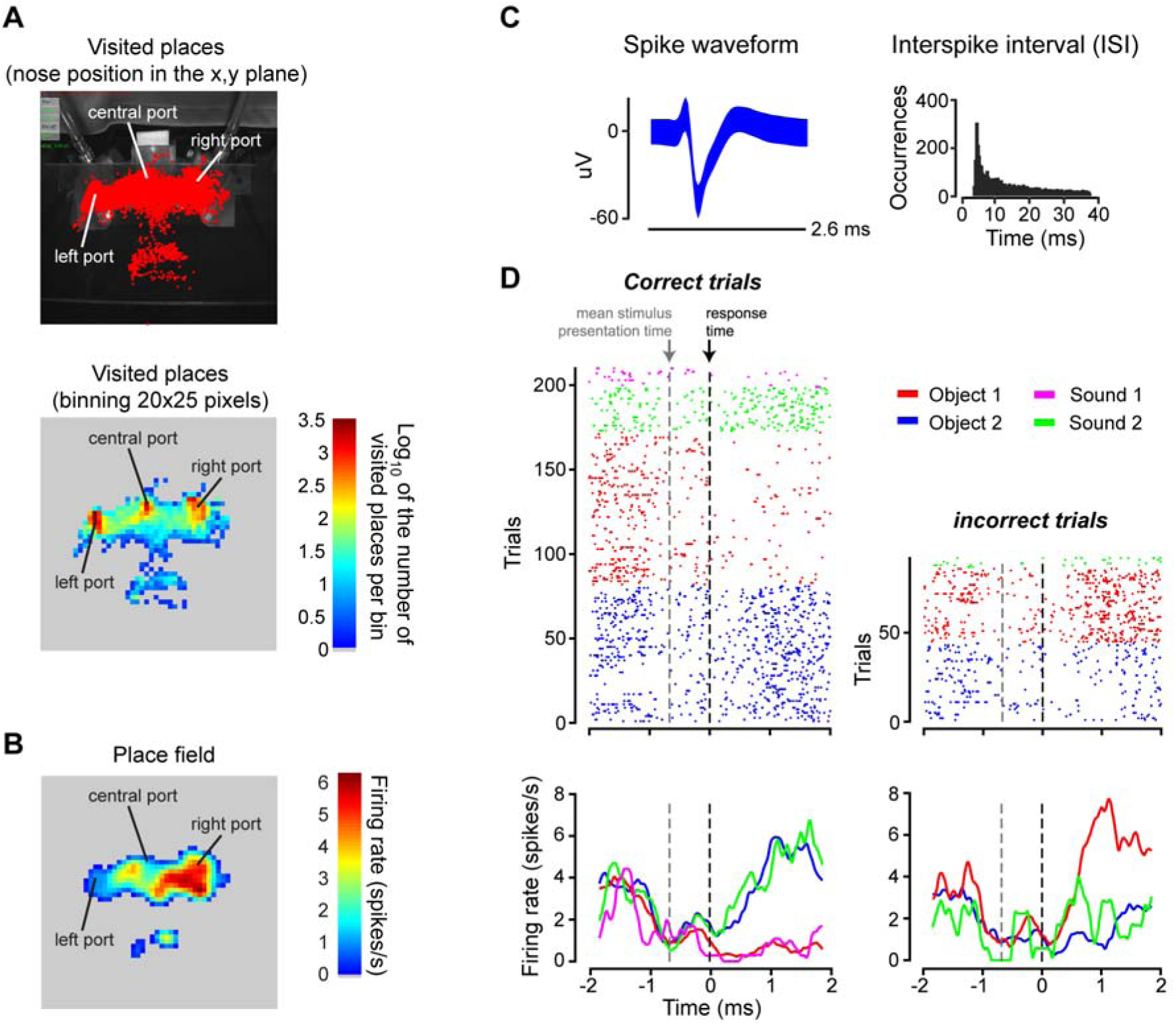
Place field of a rat hippocampal neuron during the execution of the perceptual discrimination task. (**A**) Top: map of the places visited by the nose of the rat (red dots) around the area of the response ports, while the animal was performing the discrimination task. Bottom: density map of visited locations around the response ports. The map was obtained by: 1) dividing the image plane in spatial bins of 20×25 pixels; 2) counting how many times the rat visited each bin; and 3) taking the logarithm of the resulting number of visits per bin (so as to allow a better visualization of the density map). (**B**) Place field of a well-isolated hippocampal neuron (as shown in **C**). The average firing rate of the neuron (i.e., the number of spikes fired in a 33 ms time bin, corresponding to the duration of a frame captured by the head tracker) was computed across all the visits the rat’s nose made to any given spatial bin around the response ports (same binning as in **B**). Only bins with at least 10 visits were considered, and the raw place field was smoothed with a Gaussian kernel with sigma = 1.2 bins. (**C**) Superimposed waveforms (left) and inter-spike interval distribution (right) of the recorded hippocampal neuron. (**D**) Time course of the activity of the neuron, during the execution of the task. The dots in the raster plots (top) indicate the times at which the neuron fired an action potential (or spike). The peri-response time histograms (PRTHs) shown in the bottom were obtained from the raster plots by computing the average number of spikes fired by the neuron across repeated trials in consecutive time bins of 0.48 ms. The color indicates the specific stimulus condition that was presented in a given trial (see caption in the figure). The panels on the left refer to trials in which the rat’s choice was correct, while the panels on the right refer to trials in which his response was incorrect. Both the raster plots and the PRTHs are aligned to the time of arrival to the response port (time 0; black dashed line). The gray dashed line shows the mean stimulus presentation time (gray dashed line), relative to the time of the response.

To verify that the neuron mainly coded positional information, we plotted its activity as a function of time, across the various epochs of the discrimination task, as well as a function of the sensory stimuli the rat had to discriminate and of the correctness of his choices. This is illustrated by the raster plots of Figure 9D (top), where each dot shows the time at which the neuron fired an action potential, before and after the animal made a perceptual choice (i.e., before and after he licked one of the lateral response ports). Individual rows show the firing patterns in repeated trials during the task, with the color coding the identity of the stimulus (see caption). In addition, the trials are grouped according to whether the rat’s response was correct (left) or not (right). Each raster plot was then used to obtain the peri-response time histograms (PRTHs) shown in the bottom of Figure 9D, where the average firing rate of the neuron across trials of the same kind is reported as a function of time. As expected for a place cell, and consistently with the spatial tuning shown in Figure 9B, the neuron started to respond vigorously after the rat licked the right response port and did so regardless of whether the stimulus was auditory or visual (see, respectively, the green and blue dots/lines in Figure 9D). In addition, the neuron also fired when the rat licked the right port in response to the visual stimulus (Object 1) that required a response to be delivered on the left port, i.e., on trials in which his choice was incorrect (red dots/line).

Obviously, our analysis of this example neuronal response pattern is far from being systematic. It is simply meant to provide an illustration of how our light-weight, portable head-tracking system can be applied to study the motor/behavioral patterns of small rodents (and their neuronal underpinnings) in experimental contexts where the animals do not navigate large environments but are confined to a restricted (often small) span of spatial locations.

## Materials and methods

### Experimental rig for behavioral tests and in-vivo electrophysiology

The rig used to administer to the rat the two-alternative forced-choice (2AFC) discrimination task (Zoccolan, 2015; Zoccolan and Di Filippo, 2018) has been already described in the Results. Briefly (with reference to Figure 1 A and B), a custom-made operant box was designed with the CAD software SolidWorks (Dassault Systèmes) and then built using black and transparent Plexiglas. The box was equipped with a 42-inch LCD monitor (Sharp, PN-E421) for presentation of visual stimuli and an array of three stainless steel feeding needles (Cadence Science), 10 mm apart from each other connected to three proximity sensors. The left and right feeding needles were connected to two computer-controlled syringe pumps (New Era Pump Systems NE-500), for automatic pear juice delivery. Each feeding needle was flanked by a light-emitting diode (LED) on one side and a photodiode on the other side, so that, when the rat licked the needle, he broke the light bean extending from the LED to the photodiode and his responses was recorded. The front wall of the operant box had a rectangular aperture, which was 4-cm wide and extended vertically for the whole height of the wall, so as to allow room for the cables connecting the implanted electrode array to the preamplifiers of the acquisition system. The rat learned to insert his head through this aperture, so as to reach the feeding needles and face the stimulus display, which was located 30 cm apart from his nose. Two speakers positioned at the sides of the monitor were used for playing the sound stimuli. Stimulus presentation, input and output devices, as well as all task-relevant parameters and behavioral data acquisition were controlled with the freeware, open-source software MWorks (https://mworks.github.io/) running on a Mac Mini (Apple; solid cyan and black arrows in Figure 1A).

During the neurophysiological experiments, stimulus presentation and collection of behavioral responses were synchronized with the amplification and acquisition of extracellular neuronal signals from hippocampus (red arrow in Figure 1A), performed using a system three workstation (Tucker-Davis Technologies – TDT), with a sampling rate of 25 kHz, running on a Dell PC. Specifically, MWorks sent a code with the identity and time of the stimulus presented in every trial to the TDT system via a UDP connection (dotted black arrow). The TDT system also controlled the acquisition of the frames by the camera of the head tracker, by generating a square wave that triggered the camera image uptake (dashed black arrow). The camera, in turn, generated a unique identification code for every acquired image. This code was saved by the PC running the head-tracking software (along with the Cartesian coordinates and pose of the head) and also fed back to the TDT (dashed green arrow), which saved it in a data file along with the neurophysiological and behavioral recordings. The square wave had a fixed period and duty cycle, but it was adjustable by the user from the TDT graphical interface. In our experiments, the period was 23 ms (about 50 Hz) and the duty cycle was around 33%.

### Perceptual discrimination tasks, surgery and neuronal recordings

One adult male Long Evans rat (Charles rivers Laboratories) was used for the validation of the head tracking system. The animal was housed in a ventilated cabinet (temperature controlled) and maintained on a 10/14-h light/dark cycle. The rat weighed approximately 250 gr at the onset of the experiment and grew to over 500 gr at the end of the study. The rat received a solution of water and pear juice (ratio 1:5) as a reward during each training session and, in addition, he had access to water ad libitum for 1 h after the training. All animal procedures were in agreement with international and institutional standards for the care and use of animals in research and were approved by the Italian Ministry of Health: project N. DGSAF 22791-A, submitted on Sep. 7, 2015 and approved on Dec. 10, 2015 (approval N. 1254/ 2015-PR).

The rat was trained in a visual and sound recognition task in the rig described in the previous section. The task required the animal to discriminate between either two visual objects or two sounds. Each trial started when the animal touched the central response port, which triggered the stimulus presentation. After the presentation of the stimulus on the monitor or the playback of the sound, the rat had to report its identity by licking either the left or the right response port. Each stimulus was associated to one specific port. Hence, only one action, either licking the left or right port, was associated to the reward in any given trial. Correct object identification was followed by the delivery of the pear juice-water solution, while incorrect response yielded a 1-3 s time out, with no reward delivery and a failure tone played along with the flickering of the monitor from black to middle grey at 15 Hz. The stimuli were presented for 1 sec or until the animal licked one of the lateral response ports, independently of the correctness of the choice.

The visual stimuli consisted of a pair of three-lobed objects, previously used by our group in several studies of visual perceptual discrimination in rats (Zoccolan et al., 2009; Alemi-Neissi et al., 2013a; Rosselli et al., 2015a; Djurdjevic et al., 2018). Specifically, each object was a rendering of a three-dimensional model built using the ray tracer POV-Ray (http://www.povray.org). The sound stimuli were two pure tones, at 600 Hz and a 6000 Hz respectively (sound level: 55 dB approximately). Sounds were delivered from two speakers located symmetrically on the both sides of the front part of the operant box, so that the sound level at the position of the animal’s ears was equal when the animal’s nose was near the central response port at the onset of the trial.

Once rat reached ≥ 70 % correct behavioral performance, it was implanted with an electrode array for chronic recordings. To this aim, the animal was anaesthetized with isofluorane and positioned in a stereotaxic apparatus (Narishige, SR-5R). A craniotomy was made above the dorsal hippocampus and a 32-channel Zif-clip array (Tucker-Davis Technologies Inc., TDT) was lowered into the craniotomy. Five stainless steel screws were inserted into the skull (three anterior, one lateral and two posterior to the craniotomy) in order to give anchoring to the implant cementation. Around the implant, we put Haemostatic gelatin sponge (Spongostan™ dental, Ethicon, Inc) saturated with sterile sodium chloride solution to protect the brain from dehydration, and then silicon (Kwik-Cast™, World Precision Inst) to seal the craniotomy and protect it from the dental cement (Secure, Sun Medical Co LTD) that was finally used to secure the whole implant to the skull. Hippocampal stereotaxic coordinates were: −3.7 mm AP, −3.5 mm ML. The final depth of the electrodes to target CA1 was around −2.2mm, and for the CA3 subregion was around −3.4mm. In order to place over the head the rat the dots’ pattern that was necessary for head tracking (see next sections), a complementary connector to the one mounted on the pattern was also cemented on the head, anterior to the electrode array (with a magnet on top; 210 g/cm^2^ holding force; Figure 2A-B). The rat was given antibiotic enrofloxacin (Baytril; 5 mg/kg) and caprofen (Rimadyl; 2.5 mg/kg, subcutaneous injection) for prophylaxis against infections and postoperative analgesia for the next three days post-surgery. The animal was allowed to recover for 7 to 10 days after the surgery, during which he had access to water and food ad libitum. The behavioral and recording sessions in the operant box were resumed after this recovery period. Action potentials (spikes) in the extracellular signals acquired by the TDT system were detected and sorted for each recording site separately, using Wave Clus (Quiroga et al., 2004) in Matlab (The MathWorks). Online visual inspection of prominent theta waveforms in addition to histology confirmed the position of the electrodes.

### Head-tracking system

The head tracker (Figure 1) consists of an industrial monochromatic CMOS camera (Point gray FLEA model), a far-red illuminator, a dedicated PC (Intel Xeon HP Workstation Z620 with Xeon CPU 2.5GHz, and RAM 16GB), and a three-dimensional pattern of dots, mounted over the head of the animal and imaged by the overhead camera (Figure 2A). The illuminator was made of a matrix of 4×4 LEDs, with dominant wavelength at 730 nm (OSLON SSL 150, PowerCluster LED Arrays) and a radiance angle of [−40°,40°], and was powered at 100-150 mW. It was mounted right above the stimulus display, oriented towards the operant box with an angle of approximately 50° with respect to the display (Figure 1B). The camera was set in the external trigger mode and the triggers it received from the TDT system (dashed black arrow in Figure 1A) were counted, so that each frame was assigned a unique code, which was encoded in the first four pixels of each acquired image. The same codes were sent to the TDT (dashed green arrow), which stored them, so as to allow the image frames (and, therefore, the positional and pose information returned by the head tracker) to be aligned to the neurophysiological and behavioral data. In our validation analyses and in our tests with the implanted rat, the CMOS sensor integration time was set at 3 ms. This value was chosen after preliminary tests with the implanted rat performing the discrimination task, in such a way to guarantee that the images of the dots’ pattern acquired during fast sweeps among the response ports were not blurred. The camera was placed above the stimulus display, oriented towards the operant box with an angle of approximately 50° with respect to the display (Figure 1B), although some of the validation measures presented in the Results (Figure 3) were obtained with the camera mounted in such a way to have its optical axis perpendicular to the floor of the measurements’ area.

Obviously, following the generation of a trigger signal by the TDT, a propagation delay occurred before the camera started integrating for the duration of the exposure time. This delay was about 5 µsec and has been measured as suggested by the camera constructor. That is, we configured one of the camera’s GPIO pins to output a strobe pulse and we connected to an oscilloscope both the input trigger pin and the output strobe pin. The time delay from the trigger input to the complete image formation was then given by the sum of the propagation delay and exposure time, where the latter, as explained above, was fixed at 3 ms. Any other possible time delays like the sensor readout and the data transfer toward the PC, do not affect real-time acquisition, since the content of the image and its frame number is entirely established at the end of the exposure time

After the image has been read from the camera memory buffer, it becomes available to the head-tracking software (see next section) and, before any image processing occurs, the embedded frame number is extracted from the image and sent back to the TDT system via UDP protocol. Due to the data transfer over the USB 3.0 and UDP connections, the frame number information reaches the TDT system with an unpredictable time distance from the signal that triggered the acquisition of the frame itself. However, synchronizing the triggering pulses and the frame-numbers is straightforward when considering the frame numbers and the number of trigger pulses generated. To validate our synchronization method, we conducted a double check by comparing the synch by frame-number with the synch by timestamps. The second method uses the timestamp of the frame-number received by TDT and looks for the trigger pulse that could have generated the frame in an interval of time [−0.0855 −0.0170] msec before the frame number arrived. In particular, it assigns to the current frame-number the more recent pulse that has not yet been assigned before. The two methods gave identical results over long sessions.

As explained in the Results, the head tracker works by imaging a 3D pattern (shown in Figure 2A), consisting of 5 coplanar black dots over a white background (approximately 3.5 mm apart from each other), plus a sixth dot located over an elevated pillar (approximately 5 mm tall). The 5 coplanar dots were arranged in a L-like shape, while the elevated dot was placed at the opposite side with respect to the corner of the L shape, so as to minimize the chance of occluding the other dots. The 3D structure holding the dots was designed with the CAD software SolidWorks (Dassault Systèmes) and then 3D printed. The dots were printed on a standard white paper sheet, which was then cut to the proper size and glued over the 3D printed structure. This structure also included an arm, with a magnet at the bottom, which allowed the pattern to be mounted over an apposite support (holding a complementary magnet) that was surgically implanted on the rat’s head (Figure 2A-B).

Since the head tracking algorithm (see next sections) requires a precise knowledge of the spatial arrangement of the dots relative to each other, the distance between each pair of dots, as well as the height of the pillar, were carefully measured using a caliper and by acquiring high-magnification images of the pattern along with a precision ruler placed nearby (SE® resolution 0.5 mm). This information is inserted in an apposite configuration file and is part of the calibration data that are required to operate the head tracker. The other calibration information is the internal camera parameters *K*, which have to be pre-computed by the camera calibration procedure (see apposite section below). Once this information is known, it can be stored and then loaded from the configuration file at the beginning of each behavioral/recording session, provided that the position of the camera in the rig is not changed and the same dots’ pattern is always used.

### Overview of the head-tracking algorithm

Our head-tracking algorithm consists of three different functional parts: 1) the *Point Detection* (PD) module; 2) the *Points-correspondences Identification* (PcI) module; and 3) the *Perspective n Point* (PnP) module. These modules are executed in a sequence at each frame captured by the camera. An estimate of the position/pose of the dot’s pattern (and, therefore, of the head) is computed, only when the PD module is able to extract the positions of all the dots within the acquired image. In cases in which such condition is not satisfied, the PD algorithm signals and records the inability to successfully recover the position/pose of the pattern for the current frame.

The PD module is a fast feature-detection algorithm that eliminates false positives blobs and restricts the search space of point configurations by certain geometric constraints. Following the correct extraction of the dots’ positions by the PD algorithm, such positions must be univocally matched to the known geometry of the pattern by the PcI algorithm. That is, all six points, as they appear in the image, must be associated to the known 3D coordinates of the corresponding dots on the 3D pattern. Finally, the position and pose of the *pattern reference system* (*x*’, *y*’, *z*’; purple arrows in Figure 2C) with respect to the *camera reference system* (*x, y, z*; black arrows in Figure 2C) is obtained by solving the PnP problem. The whole algorithm was designed to process the images captured by the camera in real-time, so as to output the estimated position and pose of the pattern without the need of storing the images for off-line processing. This required designing the three different modules, in such a way to maximize both the speed of processing and the accuracy of the estimates.

### Point detection (PD) module

To solve the final PnP problem and estimate the pose of the dots’ pattern in the camera reference system, all the possible point candidates that represent a 2D projection configuration of the pattern must be considered. This requires extracting first the positions of all six points in each frame captured by the camera. The PD module of our algorithm takes care of this step by applying a Difference of Gaussians (DoG) filter, which is particularly efficient to compute, is rotationally invariant, and shows a good stability under projective transformation and illumination changes (Lowe, 1991, 2004).

The DoG filter is an approximation of the well-known Laplacian of Gaussian (LoG) filter. It is defined as the difference between the images resulting from filtering a given input image *I* with two Gaussians having different sigma, *σ*_1_ and *σ*_2_ (Jähne, 2005), i.e.: DoG = G(*I,σ*_1_) – G(*I,σ*_2_), where G(*I,σ*) is the convolution of *I* with a Gaussian filter *G* with parameter *σ*. When the size of the DoG kernel matches the size of a blob-like structure in the image, the response of the filter becomes maximal. The DoG kernel can therefore be interpreted as a matching filter (Duda et al., 2001). In our implementation, the ratio *R* = *σ*_1_ /*σ*_2_ has been fixed to 2. Therefore, *σ*_1_ and *σ*_2_ can be written as:

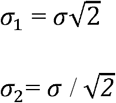

where *σ* can be interpreted as the size of the DoG kernel. In principle, *σ* should be set around 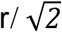, where *r* is the radius of a black dot as it appears on a frame imaged by the camera (Lindeberg, 1998). However, the method is quite tolerant to the variations of the distance of the dot’s pattern from the camera, and the dots are correctly detected even when they are at a working distance that is half than that originally set (i.e., when their size is twice as large as *r*). In addition, the software implementation of our head tracker includes a GUI that allows manually adjusting the value of σ, as well as of other key parameters (e.g., *Ic* and *score;* see next paragraph/sections), depending on the stability of the tracking procedure (as visually assessed by the user, in real-time, through the GUI).

After detecting the candidate dots in the image plane using the DoG filter, we applied a *non-maxima suppression* algorithm (Canny, 1987) that rejects all candidate locations that are not local maxima and are smaller than a contrast threshold *Ic*. Still, depending on the value of *Ic*, a large number of false positives can be found along the edges of the pattern. In fact, a common drawback of the DoG and LoG representations is that local maxima can also be detected in the neighborhood of contours or straight edges, where the signal change is only in one direction. These maxima, however, are less stable, because their localization is more sensitive to noise or small changes in neighboring texture. A way to solve the problem of these false detections would be to analyze simultaneously the trace and the determinant of the Hessian matrix over a neighborhood of pixels in the image (Mikolajczyk and Schmid, 2002). The trace of the Hessian matrix is equal to the LoG but considering simultaneously the maxima of the determinant penalizes points for which the second derivatives detect signal-changes in only one direction. Since a similar idea is explored in the *Harris cornerness* operator (Harris and Stephens, 1988), our algorithm exploits the already calculated Gaussian smoothed image and uses the efficient implementation of Harris corner detection available in the OpenCV library (https://opencv.org).

Given an image *I*, Harris cornerness operator obtains the local image structure tensor 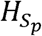 over a neighborhood of pixels *S*_*p*_, where 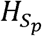 is defined as:

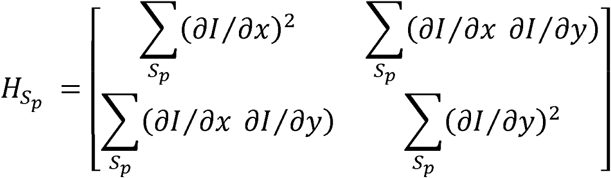

Here, ∂*I*/ ∂*x*,and, ∂*I*/ ∂*y* are the partial derivatives of the image intensity *I* along the two spatial axes *x* and *y* of the image plane, computed using a Sobel operator with an aperture of 3 pixels – i.e., a 3×3 filter that implements a smooth, discrete approximation of a first order derivative (Jähne, 2005; González and Woods, 2008). 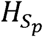 provides a robust distinction between edges and small blobs because the difference det 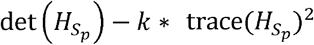 assumes values that are strictly negative on edges and positives on blob centers. This difference depends on three parameters: 1) the aperture of the Sobel filter, which, as mentioned above, was fixed to the minimal possible value (3) for the sake of speed of computation; 2) the weight *k*, assigned to the trace of the 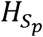 tensor, which, as suggested in (Grauman and Leibe, 2011), was set to 0.04, i.e., the default value of the openCV library (https://opencv.org); and 3) the block size of the squared neighborhood *S*_*p*_ that was empirically set to 9 based on some pilot tests, where it showed good stability over a broad range of working distances (note that *S*_*p*_ could be extrapolated from the dots’ size, the working distance between the camera and the dots’ pattern and the internal camera parameters, assuming an ideal condition of a clean white background around the dots).

To summarize, the complete PD algorithm worked as follows. First, the DoG filter was applied to identify all candidate dots in the acquired image. Then, the non-maxima suppression algorithm was used to prune some of the false positives around the maxima. Finally, for each of the remaining detected dots, the Harris cornerness operator 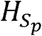 was computed over the a neighborhood *S*_*p*_ centered on the position of each dot and, depending on the sign of the difference det 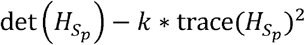, the dot was rejected as a false positive or accepted as the projection on the image plane of one of the dots of the 3D pattern. As mentioned above, the contrast threshold *Ic* of the non-maxima suppression algorithm was adjustable by the user through the same GUI used to adjust *σ* of the DoG.

To conclude, it should be noted that, in spite of the pruning of false detections performed by the PD algorithm, still many spurious dots are identified in the image, in addition to those actually present on the pattern. This is because, in general, many dot-like features are present in any image. To further refine the identification of the actual dots of the pattern, it is necessary to take into account their relationship, given the geometry of the pattern itself. This is achieved by the PcI algorithm that is described in the next section.

### Point-correspondences identification (PcI) module

Since projective transformations maintain straight lines, aligned triplets of dots (p1, p2, p3) in the 3D pattern must still be aligned in the images captured by the camera. Our PcI algorithm searches all aligned triplets of dots in an image and, in order to reduce the number of possible triplets, only those having length (i.e., distance between the external points) smaller than *D* are considered. To identify a triplet, for any given pair of detected dots, we looked whether a third, central dot was present in the proximity of the middle position in-between the pair (note that, in case of a triplet, the offset of the projected central point with respect to the middle position is negligible, because usually the black dots are at a much shorter distance than the working distance and the orthographic projection can be adopted). *D* is automatically computed from the physical distances of the external points of the triplets on the dot’s pattern, knowing the intrinsic parameters of the camera. It must be set to be slightly bigger (10% bigger showed to be widely sufficient) than the maximum distance between the external points in the triplets on the image plane, when the pattern is positioned at the maximal working distance from the camera. This condition is achieved when the z-axis of the pattern is exactly aligned to the optical axis of the camera.

Once all candidate triplets have been identified, the algorithm looks for those having a common external point, which corresponds to the corner of the L-shaped arrangement of 5 coplanar dots in the pattern (Figure 2A). The search of this L’s corner is performed by considering 5-tuples of points configurations to obtain the correspondence final assignment. In addition to collinearity, another important projective invariant is the angular ordering (on planes facing the view direction of the imaging device). That is, if we take three points defining a triangle, once we have established an ordering to them (either clockwise or anti-clockwise), such ordering is maintained under any projective transformations that looks down to the same side of the plane (Bergamasco et al., 2011). In our framework, this implies evaluating the external product of the two vectors that start from the common point (i.e., the L’s corner) and end on the respective external points. This establishes the orientation order and, consequently, assigns uniquely the 5 black dots correspondences between the 3D pattern and its image.

Following this assignment, the 6th dot belonging to the pattern (i.e., the one placed over the pillar; Figure 2A) is searched in proximity of L’s corner. To this aim, the maximal distance between the dot on the pillar and the L’s corner (*DP*) in the image plane is automatically estimated (in the same way as *D*) from the actual distance between the two dots in the 3D pattern, knowing the camera’s internal parameters and its maximal working distance from the pattern. This yields a set of candidate pillar dots. Finding the correct one requires evaluating each candidate dot in conjunction with the other 5 coplanar dots in terms of its ability to minimize the *reprojection error* computed by solving the PnP problem (see next section). It should be noticed that the correct identification of the 6^th^ point on the pillar is fundamental, since it is the only point out of the plane that allows the PnP problem to be solved robustly. The reprojection error is defined as the sum norm distance between the estimated positions of the dots in the image and the projections of the physical dots of the 3D pattern on the image plane, under a given assumed pose of the pattern and knowing the camera’s calibration parameters. More specifically, during the 6^th^ point selection we defined the score of a given dots’ configuration as:

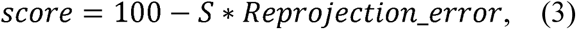

where *S* is a proper scalar factor established experimentally (in our application, we set *S* = 5). To understand the meaning of *S*, let’s suppose, for instance, to have a distance error of one pixel for each dot, thus yielding a reprojection error of 6. Without the *S* scaling factor (i.e., with *S* = 1), we would obtain a *score* of 94. However, the error on each dot is typically well below one pixel (see next paragraph about the way to estimate the dots’ coordinates with sub-pixel accuracy) and the score would therefore be always close to 100. Hence, the need of introducing the factor *S* to rescale the score, so that it can range between 90 and 100. As mentioned above, during the selection procedure, the 6^th^ point on the pillar was chosen among the candidates that maximized the score defined above. In fact, each hypothesis about the position of the 6^th^ point yielded a PnP transformation, for which it was possible to compute the reprojection error. Note that, to eliminate some possible ambiguities in the selection of the 6^th^ point, we also exploited the a-priori knowledge about the direction of the pillar (which must point toward the camera sensor).

As expected, given how sensitive the PnP procedure is to small variations in the estimated positions of the dots, we empirically verified that a pixel-level accuracy was not sufficient to guarantee high precision and stability in our pose estimates. For this reason, we estimated the positions of the centers of the dots at the sub-pixel level by 3-point Gaussian approximation (Naidu and Fisher, 1991). This method considers the three highest, contiguous intensity values (along either the *x* or *y* spatial axes) within a region of the image that has been identified as one of the dots (i.e., a blob) and assumes that the shape of the observed peak fits a Gaussian profile. This assumption is reasonable, because the sensor integration over a small area of pixels containing a blob, after the DoG filtering, produces smooth profiles very similar to Gaussian profiles. If *a, b* and *c* are the intensity values observed at pixel positions *x* — 1, *x* and *x* + 1 with *b* having the highest value, then the sub-pixel location (*x*_*s*_) of the peak is given by:

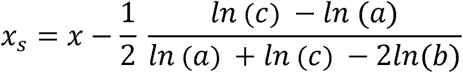

where *x* is the x-coordinate of the center of the pixel with intensity value *b*. The same approximation is applied to obtain the sub-pixel *y* coordinate *y*_*s*_ of the dot center.

### Perspective n Point (PnP) module

In computer vision, the problem of estimating the position and orientation of an object with respect to a perspective camera, given its intrinsic parameters obtained from calibration and a set of world-to-image correspondences, is known as the Perspective-n-Point camera pose problem (PnP) (Lowe, 1991b; Fiore, 2001). Given a number of 2D-3D point correspondences *m*_*i*_*↔M*_*i*_ (where *m_i_*=[*u v*]’ are the 2D coordinates of point *i* over the image plane, and *M*_*i*_ = [*XYZ*]’ are the 3D coordinates of point *i* in the physical environment) and the matrix *K* with the intrinsic camera parameters (see definition below), the PnP problem requires to find: 1) a rotation matrix *R* that defines the orientation of the object (i.e., of the pattern reference system *x*’, *y*’, *z*’ in our case) with respect to the camera reference system *x, y, z* (see eq. 1 and Figure 2C); and 2) a translation vector *t* that specifies the Cartesian coordinates of the center of the object (i.e., of the origin *O*’ of the pattern reference system in our case) in the camera reference system, such that:

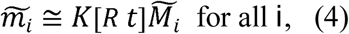

where ~ denotes homogeneous coordinates (Jähne, 2005) and ≅ defines an equation up to a scale factor. Specifically:

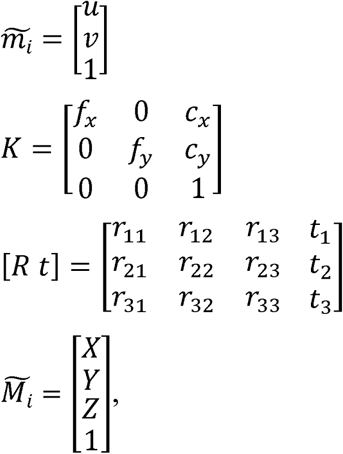

where *f*_*x*_ and *f*_*y*_ are the focal lengths and *c*_*x*_ and *c*_*y*_ are the coordinates of the principal point of the camera lens.

To solve the PnP problem, all methods have to face a trade-off between speed and accuracy. Direct methods, such as Direct Linear Transform (DLT), find a solution to a system of linear equations derived from (4) and are usually faster but less accurate, as compared to iterative methods. On the other hand, iterative methods that explicitly minimize a meaningful geometric error, such as the reprojection error, are more accurate but slower (Garro et al., 2012). In our application, we adopted the method known as EPnP (Lepetit et al., 2008), followed by an iterative refinement. The EPnP approach is based on a non-iterative solution to the PnP problem and its computational complexity grows linearly with *n*, where *n* is the number of point correspondences. The method is applicable for all *n* ≥ 4 and properly handles both planar and non-planar configurations. The central idea is to express the *n* 3D points as a weighted sum of four virtual control points. Since in our setup high precision is required, the output of the closed-form solution given by the EPnP was used to initialize an iterative Levenberg-Marquardt scheme (More, 1977), which finds the pose that minimizes the reprojection error, thus improving the accuracy with negligible amount of additional time. Both the EPnP and the Levemberg-Marquardt iterative scheme are available in the openCV library (https://opencv.org) and are extremely efficient. The execution time in our HP Workstation Z620 was in the order of some milliseconds.

### Camera calibration procedure

Camera calibration, or more precisely camera *resectioning*, is the process of estimating the parameters of a pinhole camera model approximating the true camera that produced a given image or a set of images. With the exception of the so-called self-calibration methods, which try to estimate the parameter by exploiting only point correspondences among images, a calibration-object having a known precise geometry is needed. In fact, self-calibration cannot usually achieve an accuracy comparable with that obtained with a calibration-object, because it needs to estimate a large number of parameters, resulting in a much harder mathematical problem (Zhang, 2000).

Much progress has been done, starting in the photogrammetry community, and more recently in the field of computer vision, in terms of developing object-based calibration methods. In general, these approaches can be classified in two major categories, based on the number of dimensions of the calibration objects: 1) 3D object-based calibration, where camera calibration is performed by observing a calibration object whose geometry in 3-D space is known with very good precision; and 2) 2D plane-based calibration, which is based on imaging a planar pattern shown at a few different orientations. In our application, we adopted this second option, because it has proven to be the best choice in most situations, given its ease of use and good accuracy. Specifically, we adopted the method of (Zhang, 2000), available in the OpenCV library (https://opencv.org), which, in its iterative process, also estimates some lens distortion coefficients (see the next section for a discussion on the distortion).

The fundamental equation to achieve the calibration is the same of the PnP problem, [i.e., equation (4)], and the iterative solution is based on minimizing the reprojection error defined in (3). However, in the case of the camera calibration, also the matrix *K* with the parameters is unknown, in addition to *R* and *t*. As such, solving the equation is, in principle, harder. However, the algorithm used to minimize the reprojection error does not need to run in real-time, since the calibration is performed before the camera is used for head tracking. In addition, the point correspondences *m_i_* ↔ *M_i_* over which the error is computed and minimized are the order of several hundreds, which makes the estimation of the parameters very robust and reliable. In our application, these points were the intersections of the 9×7 squares of a planar checkerboard (shown in Figure 4-figure supplement 1A) imaged in 15 different poses/positions. Specifically, a few snapshots were taken centrally at different distances from the camera, others spanned the image plane to sample the space where the radial distortion is more prominent, and finally (and more importantly) other snapshots were taken from different angles of orientation with respect to the image plane (Zhang, 2000)

To measure the effect of changing the calibration images over the pose estimation, we collected a set of 80 images of the calibration checkerboard at different orientations. 50 random subsamples (without replacement), each composed of 50% of the total images, were used to calibrate the system, thus yielding 50 different calibrations. In Table 1, we report the mean and standard deviation of the internal parameters and the lens distortion coefficients obtained from such calibrations. In Table 2, we report the averages and standard deviations of the radial and tangential part contributions of the distortion. Some of the parameters reported in these tables (i.e., *f_x_, f_y_, c_x_* and *c_y_*) have been defined in eq. (4), while other parameters (*k*_1_, *k*_2_, *k*_3_, *p*_1_ and *p*_2_) are distortion coefficients described in the next paragraphs.

**Table 1.** Mean values and standard deviations of the internal camera parameters and distortion coefficients. The reprojection error, namely the distance between the square’s corners in the images and their points reprojected after the calibration, is reported in the last column.

|  | **fx** | **fy** | **Cx** | **cy** | **k1** | **k2** | **p1** | **p2** | **k3** | **Rms**<br>**[pixel]** |
| --- | --- | --- | --- | --- | --- | --- | --- | --- | --- | --- |
| **mean** | 2701.7 | 2707.4 | 636.6 | 508.9 | -0.396 | 2.23 | 0.00098 | -0.0019 | -26.37 | 0.10 |
| **Std** | 2.51 | 1.98 | 4.73 | 4.29 | 0.012 | 0.78 | 0.00018 | 0.00025 | 15.50 | 0.00094 |

**Table 2.**
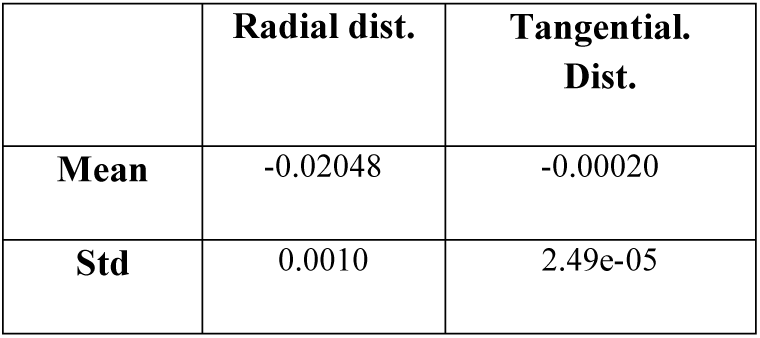
Mean values and standard deviations of the two components of the distortion, i.e. the radial and the tangential part. The radial distortion of −0.02048 indicates that the pixels in the corners of the image appear about 2% closer to the center of the image (barrel distortion).

The focal lengths *fx* and *fy* (Table 1) showed a good stability. The coordinates of the principal point (*cx, cy*), as it is well known, are some of the hardest parameters to estimate and, in our case, had a standard deviation of approximately 4.5 pixels. In any event, they are additive terms and, as such, do not affect distance and angle measurements. The remaining parameters describe the distortion introduced by the lens during the image formation. We quantified the effect of distortion, both the radial and the tangential part, on the pose estimation by considering our specific setup.

From eq. (4), after the roto-translation transformation [*x y z*] = [*R t*] [*X Y Z* 1], we have the 2d homogeneous coordinates

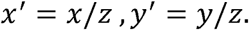

Then, the distorted coordinates *x*″ and *y*″ can be modeled by the parameters *k*_1_, *k*_2_, *k*_3_, *p*_1_ and *p*_2_ using the following equation:

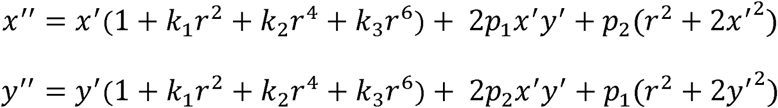

where. *r*^2^ = *x′*^2^ +*y′*^2^ The final 2D image coordinates are then obtained by:

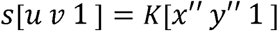

In the worst case scenario, namely in the corners of the image where the distortion is at its maximum, *x*′ and *y*′ can be approximated by *dSx*/2*f* and *dSy*/2*f*, where *f* is the focal length of the lens and *dSx* and *dSy* are the x and y dimensions of the CCD sensor. By simple geometrical considerations, since our camera mounts a 1/3” CMOS sensor and the lens has a focal length of 12.5mm, *r*^2^ results equal to 0.0576. The distortion corrections were then calculated for every sample considering this worst case. The radial distortion correction (*k*_1_*r*^2^ + *k*_2_*r*^4^ + *k*_3_*r*^6^) dominates the tangential part (2*p*_1_*x′ y′+ p*_2_(*r*^2^ + 2*x*′^2^)) but, more importantly, it has a standard deviation of just about 0.1% (see Table 2).

In this framework, it is clear that the calibration is very stable, with parameters fluctuating around 0,1% for *fx, fy* and the distortion coefficients, whereas *cx* and *cy* do not affect the displacement or angular measurements. Vice versa, an accurate measurement of the physical distances between the dots of pattern is crucial. We verified that an error of 5% in the dot distances produces errors of 5% in the pose estimates (for example, if the distances are estimated 5% bigger than the truth, the dots’ pattern is estimated at a pose 5% more distant to the camera) and, consequently, the translation measurements are affected.

## Acknowledgments

This work was supported by a Human Frontier Science Program Grant to DZ (contract n. RGP0015/2013), a European Research Council Consolidator Grant to DZ (project n. 616803-LEARN2SEE) and a FSE POR Regione autonoma FVG Program Grant (HEaD - Higher education and development) to WV. We thank Marco Gigante for his technical support in designing and building the custom assemblies used to validate the head tracker.

