## Supplementary figures for "A passive, camera-based head-tracking system for real-time, 3D estimate of head position and orientation in rodents"

**Davide Zoccolan**

International School for Advanced Studies (SISSA)

Via Bonomea, 26534136 Trieste (TS) ITALY

### Supplementary Figures

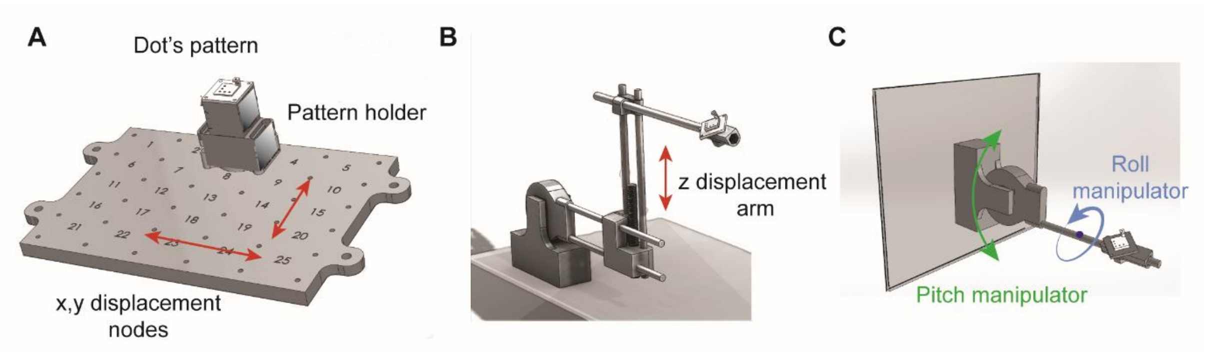


**Figure 3-figure supplement 1.** CAD drawings of the custom assemblies of breadboards, goniometers, linear and rotary stages employed to validate the head tracker. (**A**) Breadboard with the grid of 5 x 5 locations were the dots' pattern was placed to obtain the validation measurements shown in Figure 3A. The pattern was mounted over an apposite pedestal (dark gray structure) with four feet, which were inserted in matching holes over the breadboard. (**B**) The stereotax arm that was used to displace the dots' pattern vertically, to obtain the validation measurements shown in Figure 3B. (**C**) The stereotax arm that was used to change the pitch and roll angles of the dots' pattern, to obtain the validation measurements shown in Figure 3C.

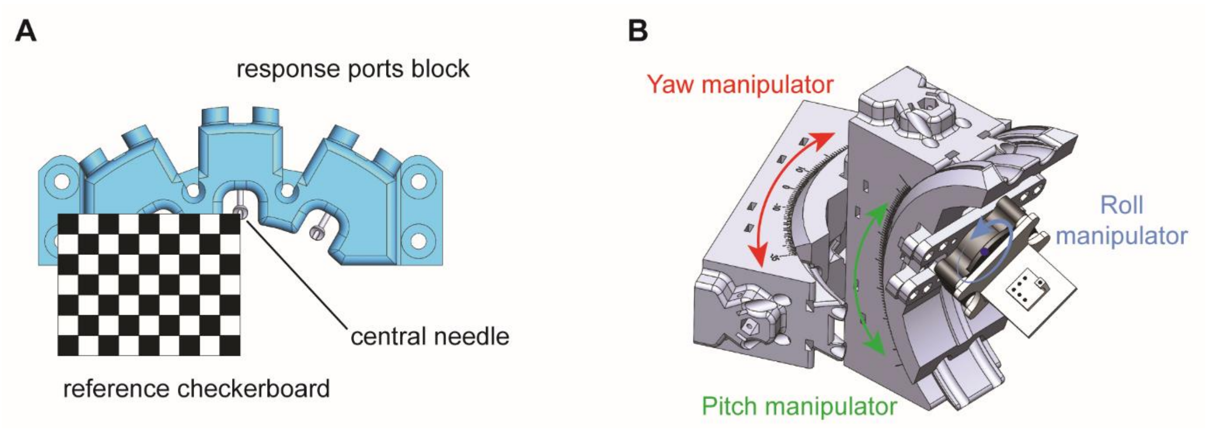

**Figure 3-figure supplement 1.** CAD drawings of the custom assemblies used to operate and validate the head tracker in a convenient reference system in the physical environment. (**A**) Top view of the 3D-printed block (cyan) holding the three feeding needles that worked as responses ports in the operant box. The figure shows how the checkerboard pattern used to calibrate the camera was placed over the response port block, in such a away to be parallel to the floor of the operant box and with its top-right vertex vertically aligned with the central needle. An image of the checkerboard pattern in such a reference position was acquired, so as to express the position measurements returned by the head tracker in a reference system with the *x* and *y* axes parallel to the edges at the base of the operant box, and the *z* perpendicular to the floor and passing through the central feeding needle. (**B**) The custom assembly of two 3D-printed goniometers and a rotary stage that was used to obtain the validation measurements shown in Figure 4B.
